# Directed Evolution of Multimodal Plastic Degrading Enzymes

**DOI:** 10.64898/2026.10.25.761710

**Authors:** Cecilia Jen Petersson, E. James Petersson

## Abstract

The inability to break down plastics is a major obstacle in the ongoing quest for sustainability. Advancements have been made in identifying enzymes that can break down certain polyesters; however, these enzymes are limited to the primary substrate and closely related plastic polymers. By increasing the catalytic promiscuity of polyester-degrading enzymes through directed evolution, a multimodal plastic degrading enzyme can be created. In order to achieve this, a polyesterase was first designed for secretion from the periplasm of *E. coli*. Through active site analysis, four sites were selected for amino acid randomization to create a >300,000 member mutant library. Next, a plate-based activity analysis system was developed to ensure efficient and reproducible screening of variants from the mutant library. Mutants with signs of activity on polyester- or polyurethane-containing agar were propagated and tested in liquid assays using a spectroscopic probe. A mutant was identified that retained polyesterase activity with newly developed polyurethane-degrading activity. Computational analysis indicated that these changes expanded the substrate pocket while retaining the positioning of the catalytic residues essential to the hydrolysis reaction. This provides the first demonstration of the directed evolution of a multimodal plastic degrading enzyme and serves as a model for further expanding polyesterase promiscuity to include multiple hydrolyzable plastics.

## Introduction

Plastic pollution is a serious threat to the health of our planet. Because plastics are cheap and easy to produce at large scales, they are often used in disposable products. For example, in 2018, 27 million tons of plastic were sent to landfills in the US alone (‘EPA - Plastics: Material-Specific Data’). This is not restricted to the US, globally 437 million tons of plastic were sold in 2022, and demand is predicted to double by 2050 given current trends (Dokl et al. 2024; Houssini, Li, and Tan 2025). Moreover, plastics in landfills take up to 500 years to decompose (Wojnowska-Baryła, Bernat, and Zaborowska 2022). Among the most common plastics are polyethylene, polypropylene, polyvinyl chloride, polystyrene, polyurethane (PU), and polyethylene terephthalate (PET), a polyester (**Figure 1A**). These plastics not only take up abundant amounts of space, but they can also find their way into the human body in the form of microplastics. Microplastics are miniature pieces of plastic that can vary from 5 mm to 1 nm,(‘EPA - Microplastics Research’) which is 80,000 times smaller than a strand of human hair. While the full extent of the effects of microplastics have not been determined yet, a variety of health conditions have been already connected to their presence in humans and animals (Huang et al. 2019; Abahussain et al. 2025). Thus, plastic pollution threatens the environment and human health, both directly and indirectly. In order to combat this growing crisis, new methods of plastic disposal must be implemented in addition to reuse and recycling approaches, which only account for ∼10% percent of plastic management (Walker 2025). Current methods being used to dispose of plastics, like incineration and landfilling, are effective for their removal from circulation, but costly for the environment and harmful to human health. For example, burning plastic can release dioxins which are toxic, cause cancer, and damage the immune system (Abahussain et al. 2025). Additionally, leaving the plastics to break down leads to the formation of microplastics which can leak out into rivers, lakes, and eventually oceans, causing widespread contamination. Scalable biodegradation methods are needed to ensure that the deterioration of plastic products is speedy, efficient, and environmentally conscious.

**Figure 1.**
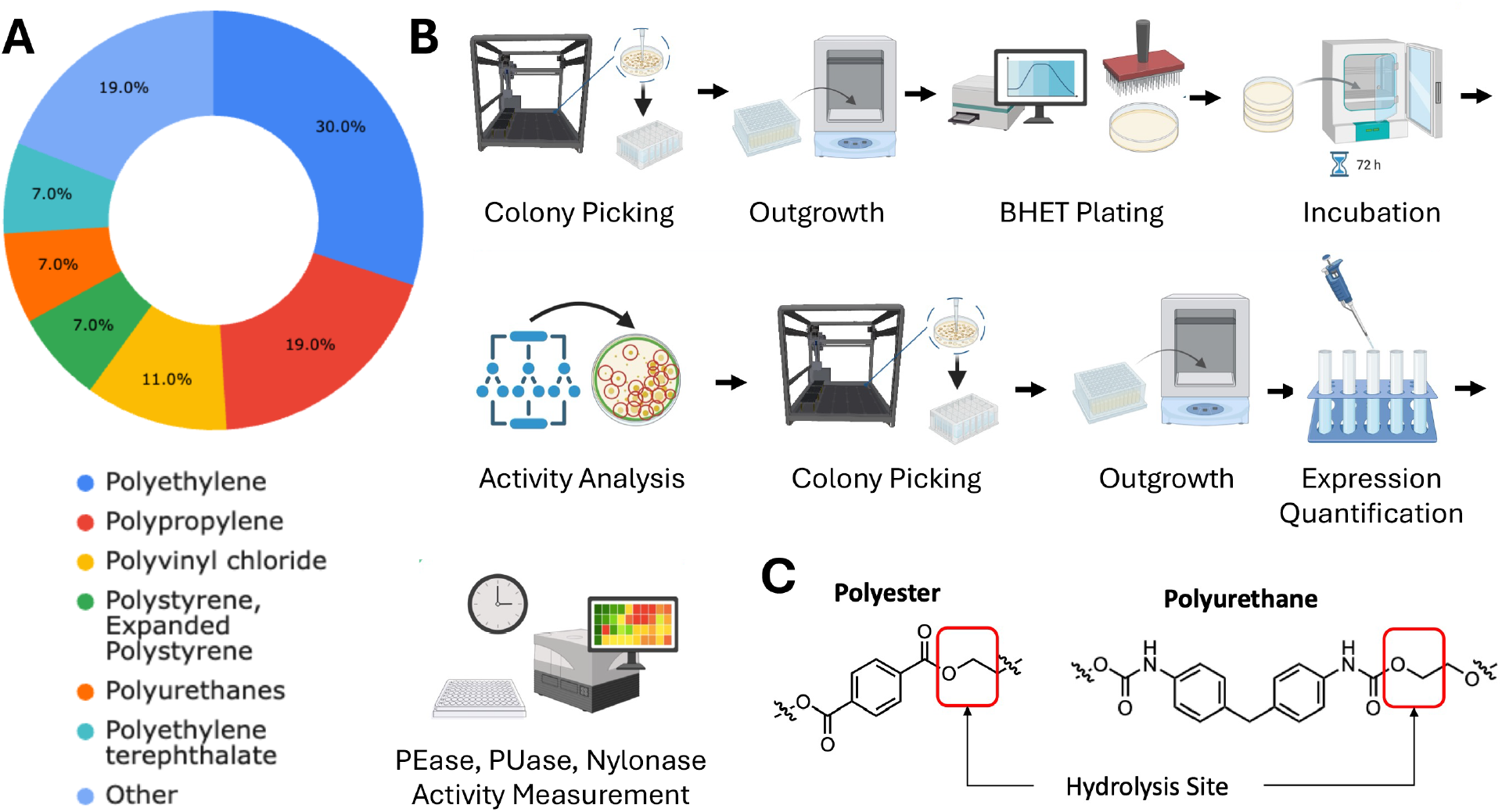
Abundance of Plastic Types and Scheme for Selection of Multi-modal Degrading Enzyme.**A.**Global plastic usage in 2018 categorized by type of resin. Data adapted from multiple sources (Zhang et al. 2022; Bertocchini and Arias 2023). **B**. Plan for design of mutant library, automated screening of PE, nylon, and PU degrading activity, and confirmation of activity in liquid assays. **C**. Hydrolyzable bonds in polyester and polyurethane. Created with BioRender.com.

Of the many plastics that are made, some can be broken down with less difficulty due to their ability to be hydrolyzed and have enzymatic degradation potential. For example, there has been successful research done on PET plastic degradation, particularly through hydrolysis of the ester bonds by PETase enzymes (Ahituv et al. 2025). Other potentially hydrolyzable groups in common plastics include the amide bonds in nylon and carbamate groups in PU. Indeed, nylonase and PUase enzymes have been identified and shown to be expressed in organisms amenable to genetic engineering and large-scale protein production like *E. coli* (Bell et al. 2024; Stern and Howard 2000). While PET only accounts for 7% of plastics, collectively, PET, nylon and PU represent about 18% of plastic pollution, comparable to polypropylene, the second most prevalent plastic after polyethylene (**Figure 1A**). Thus, a multimodal plastic hydrolyzing enzyme could address a significant portion of plastic pollution. Although mixtures of PETases, nylonases, and PUases could be used to for hydrolysis of these plastics, purified enzymes are more limited in application scale compared to enzymes secreted from bacteria for *in situ* degradation of plastics. Since maintaining cultures of three engineered bacteria each secreting a different enzyme is challenging and current nylonases are too large for secretion, we targeted engineering a secretable PETase with the ability to degrade multiple forms of plastic.

To generate a multi-modal plastic degrading enzyme, we designed a plasmid to secrete an engineered version of *Ideonella sakaiensis* (Is) PETase, called FAST-PETase and a plate-based selection system with automated activity analysis (**Figure 1B**). After optimizing the selection procedure, we generated a mutant library through saturation mutagenesis at four sites. Mutants with demonstrated activity on the initial assay plates were further cultured in liquid media and supernatants with secreted protein were tested for PUase activity in liquid culture and on PU-treated plates. Mutants with dual PETase and PUase activity identified, sequenced, and computational model of their active sites were used to analyze their multimodal hydrolysis activity.

## Results

Before engineering the desired multimodal plastic hydrolysis enzyme, we needed to develop a robust activity screening assay. To optimize growth and imaging conditions, several variants of PETase with differing levels of activity were designed for secretion to enable *in situ*, plate-based measurement of bis(2-hydroxyethyl) terephthalate (BHET) hydrolysis. Is, FAST, Hot, Thermo, and Dura PETase genes were synthesized based on previously reported sequences (plasmid map, DNA sequences, and protein sequences given in Supporting Information, SI, **Figures S1-S7**). The genes were inserted between the XbaI and XhoI sites in the pET28a(+) plasmid, placing them under control of the lac repressor for isopropyl-β-D-1-thiogalactopyranoside (IPTG) induction of protein expression. As a negative control, a Y82 to TAG mutation was made in FAST PETase to prevent translation of full length, active enzyme. All of the constructs had a pelB peptide fusion at the N-terminus to enable secretion from the periplasm. Secretion tags have previously been applied to Is PETase, and the highly similar sequences of the enzymes gave us confidence that all variants would be secreted (Cui et al. 2021; Seo et al. 2019; Shi et al. 2021). After transformation into *E. coli* BL21(DE3) cells, single colonies were picked and grown in liquid starter cultures before spotting onto BHET-containing media.

Testing of BHET clearance on solid agar media plates with kanamycin antibiotic, 1.0 mM IPTG, and 2 g/L BHET showed clear activity for several of the variants after 2 days of growth at 37 °C (SI, **Figure S8**). Various lighting, background, and camera setups were tested, and backlighting with an LED tracing tablet for uniform lighting, with the plate elevated by 10 cm above a black paper background with minimal room lighting was found to provide optimal contrast for observing clearance rings with minimal glare artifacts. Since FAST PETase showed an intermediate level of activity in our initial test, we used it to test different combinations of IPTG (0.5, or 1.0 mM) and BHET (0.5, 1.25, or 2 g/L) concentrations as well as growth temperatures (30 or 37 °C) and times (1, 2 or 3 days). We found that plates with 1.0 mM IPTG and 2 g/L BHET (hereafter referred to as BHET plates), grown for 2 days at 30 °C gave optimal consistency in activity levels between colonies. We also tested similar plate formulations using hydrolyzed, solubilized Nylon-6 and plates with PU either added to the plate media prior to pouring or top-coated after setting of the plates. Although colonies expressing FAST PETase grew, no nylon or PU clearance activity was observed (SI, **Figure S8**). Thus, our development of image analysis procedures focused on BHET clearance.

Using 2 g/L BHET solid media with incubation at 30 °C and the optimized imaging setup, clearance from 5 identical spots of each of the PETase variants was tested (**Figure 2**). No colony growth or activity was observed from Hot PETase, and Dura PETAse exhibited inconsistent growth with minimal. WT, FAST, and Thermo PETase grew consistently, with FAST and Thermo having higher activity levels than WT. Y82TAG colonies grew consistently, but had no clearance activity. Thus, we used plates with Y82TAG, WT, FAST, Dura, and Thermo to optimize our image analysis program.

**Figure 2.**
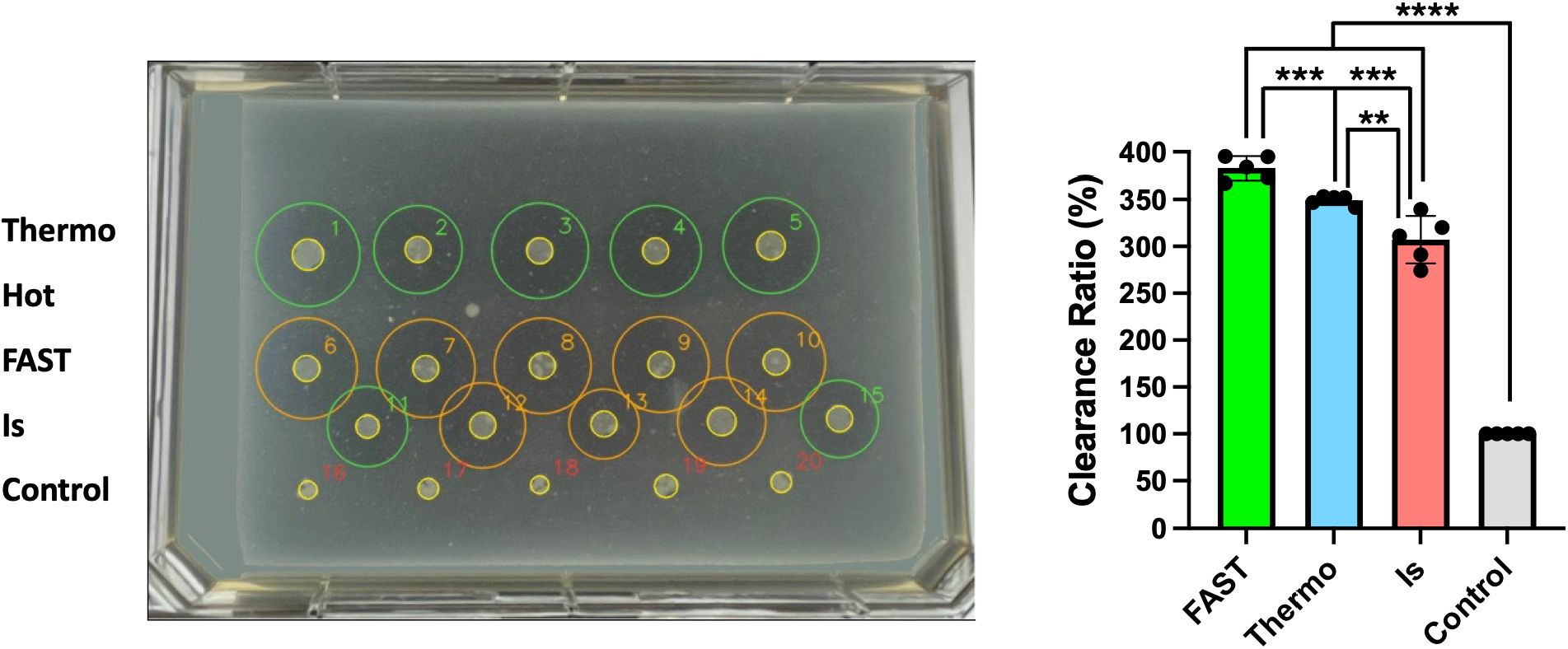
Clearance Ring Analysis.**A.**Image of BHET plate with 5 replicates each of colonies expressing Thermo, Hot, FAST, Is, and control PETases. **B**. BHET-degrading activity of each PETase variant, showing consistency of quantification. ** = p<0.01, *** = p<0.001, **** = p<0.0001

Images were collected under standardized lighting, timing, and temperature conditions. Classical image analysis was used to reliably quantify activity. Before input to the analysis module, the images were preprocessed. This included cropping to the plate region, normalizing brightness and contrast, and selecting clear agar plate regions for reference. After colonies were identified, radial intensity profiling was used to map the changes in pixel brightness to the halos around each colony. Colony activity was determined by the ratio of the halo radius to the colony radius. Classical image analysis achieved standardized results across images with clear background (for full image analysis workflow see SI, **Figure S9**). Model output was visually reviewed against raw images to confirm biologically reasonable predictions before quantitative comparisons were made.

Two FAST PETase mutant libraries were generated, the first by saturation mutagenesis at Y82, and the second by saturation mutagenesis at Y82, W151, M156, and W180. The four positions were chosen based on analysis of the active site of FAST PETase, using the crystal structure with BHET bound (Li et al. 2025). This library avoided mutation of the catalytic triad residues Ser160, His237, Asp206, instead mutating neighboring sites that formed the binding pocket for BHET in order to allow for flexible binding of related PU (carbamate) and nylon (amide) substrates (**Figure 3**).

**Figure 3.**
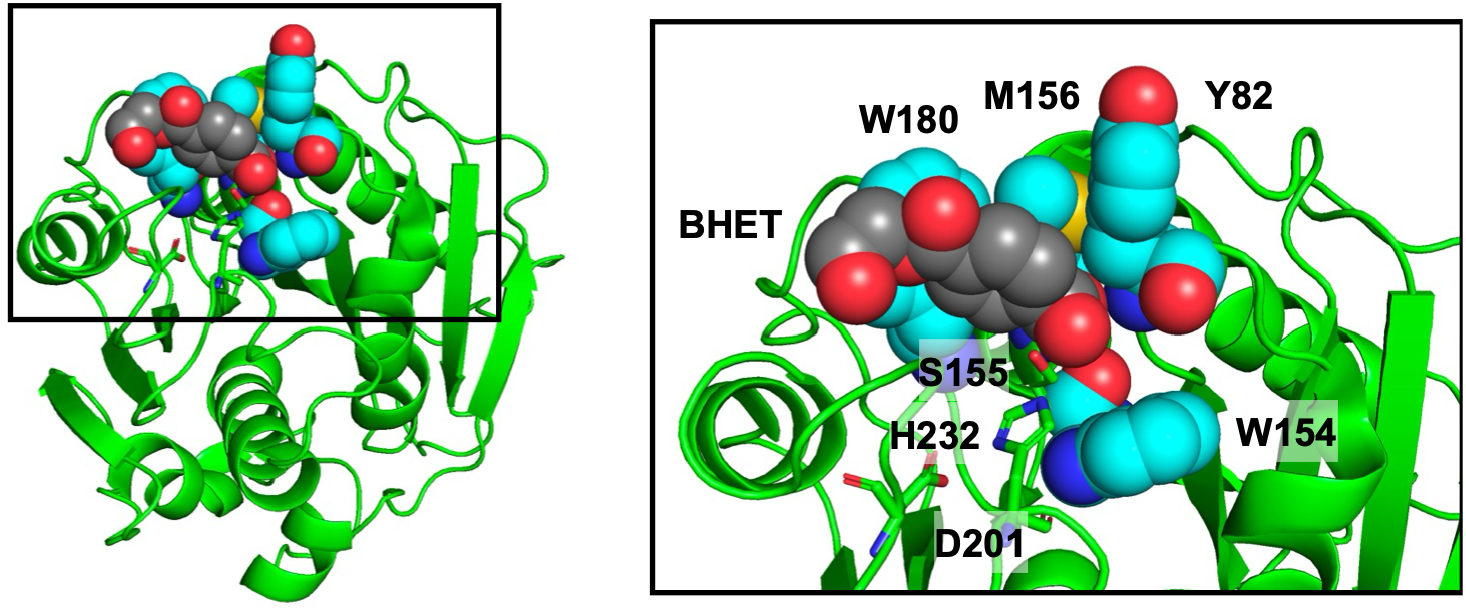
PETase Mutation Site Selection. BHET substrate and mutant library residues shown as spheres. Catalytic residues shown as sticks. Model based on PDB ID 9LMV (Li et al. 2025).

In order to maximize coverage of protein sequence space with minimal DNA libraries, the “22 codon trick” (Kille et al. 2013) was used in designing the primers for Y82, W151, and W180, with an NNK library at M156. This gave theoretical library sizes of 22 sequences for the W82 library and 340,736 sequences for the 4-position library. The W82 library was generated by standard PCR mutagenesis using the Y82TAG FAST PETase plasmid. The 4-position library was generated by PCR amplifying the 4 separate segments shown in **Figure 4**, then successive rounds of overlap extension PCR were used to combine blocks 3+4, then to combine blocks 2+(3+4), and finally to form the full length PETase mutant library insert from 1+(2+3+4). The Y82TAG FAST PETase plasmid was digested with XbaI and XhoI, then gel purified. The Y82TAG version was used to ensure that no FAST PETase background activity would be present after assembly of the mutant library plasmid. A Gibson assembly reaction was performed with the PETase mutant library insert and digested Y82TAG PETase plasmid.

**Figure 4.**
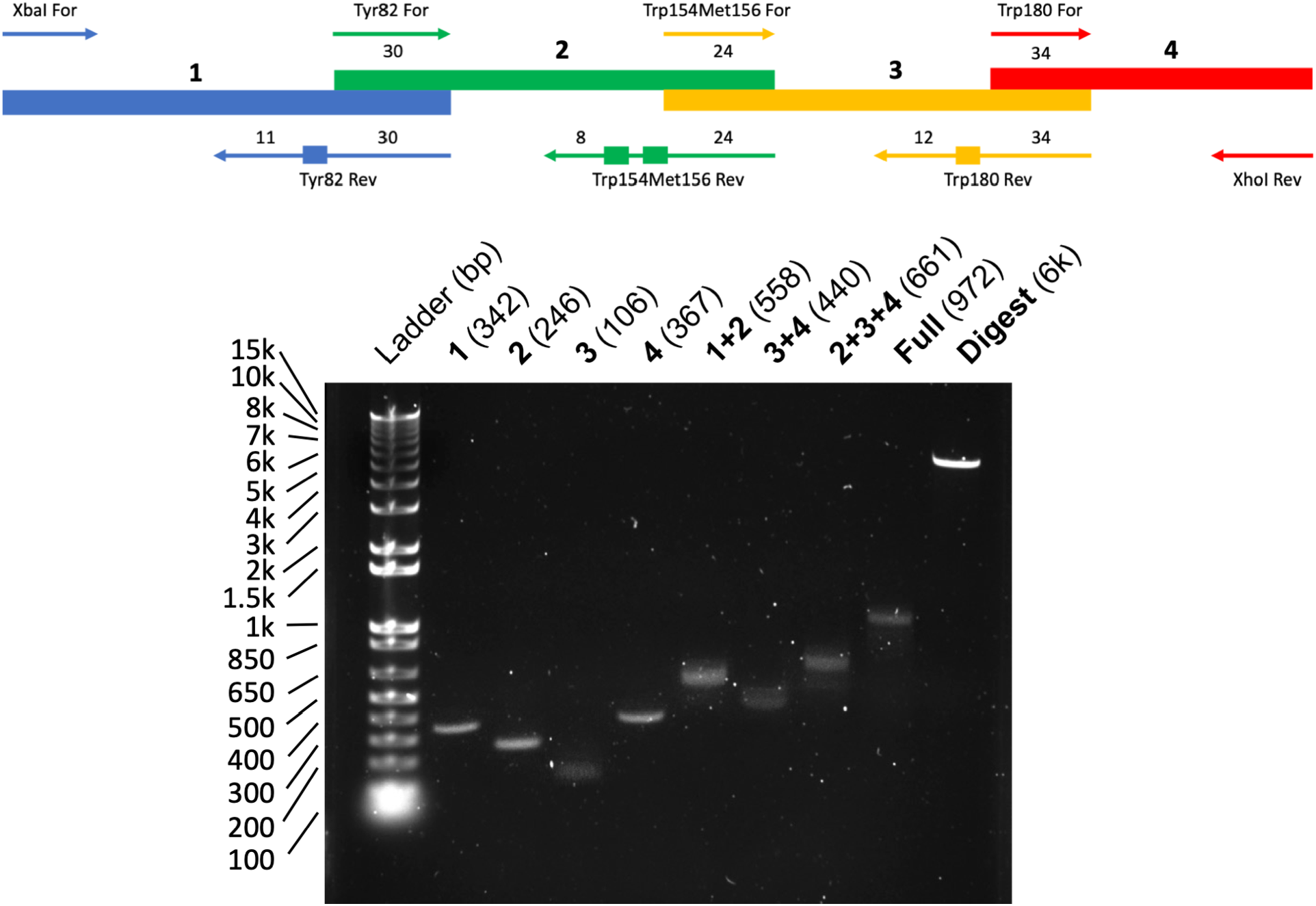
PETase Library Construction. Top: Scheme for assembly of mutant library insert by PCR amplification. Bottom: Gel-purified PCR products from amplification of segments 1, 2, 3, and 4, and then generation of combined segments 1+2, 3+4, 2+3+4, and 1+2+3+4, forming the full PETase gene (Full) are shown with XbaI/XhoI-digested Y82TAG FAST PETase PET28a plasmid (Digest). Full and digest were combined in Gibson Assembly reactions to generate the PETase mutant library.

The Gibson assembly product was transformed into *E. coli* BL21(DE3) cells by electroporation and plated on LB+Kan plates. Each transformation yielded up to ∼200 colonies. Single colonies were picked and grown in non-inducing liquid media to OD^600^ ∼0.5 in a 96 well block. These cultures were used to inoculate BHET and PU plates with a 48-pin replicator and the plates were incubated for 2 days at 30 °C for imaging (see plate images in SI, **Figure S10**).

Since the majority of mutations were expected to compromise BHET hydrolysis activity, we chose to select all colonies that showed BHET activity above background and investigate their potential for PU hydrolysis. Colonies were picked either manually or using image analysis, then cultured overnight in non-inducing LB+Kan media before plating on BHET plates to confirm activity, whole plasmid sequencing, and generation of glycerol stocks for long-term storage. From >1000 colonies in the primary BHET plate screen, 23 colonies were selected for secondary screening. All of the colonies showed some BHET clearance activity (**Figure 5**). Some of the colonies were found to be WT FAST PETAse or mixtures of sequences (see SI **Figures S11-S12** for sequencing information). Plasmids for a selection of 10 of the most active, novel PETase mutants were amplified and taken forward to testing with chromophore-based ester and carbamate (urethane) substrates.

**Figure 5.**
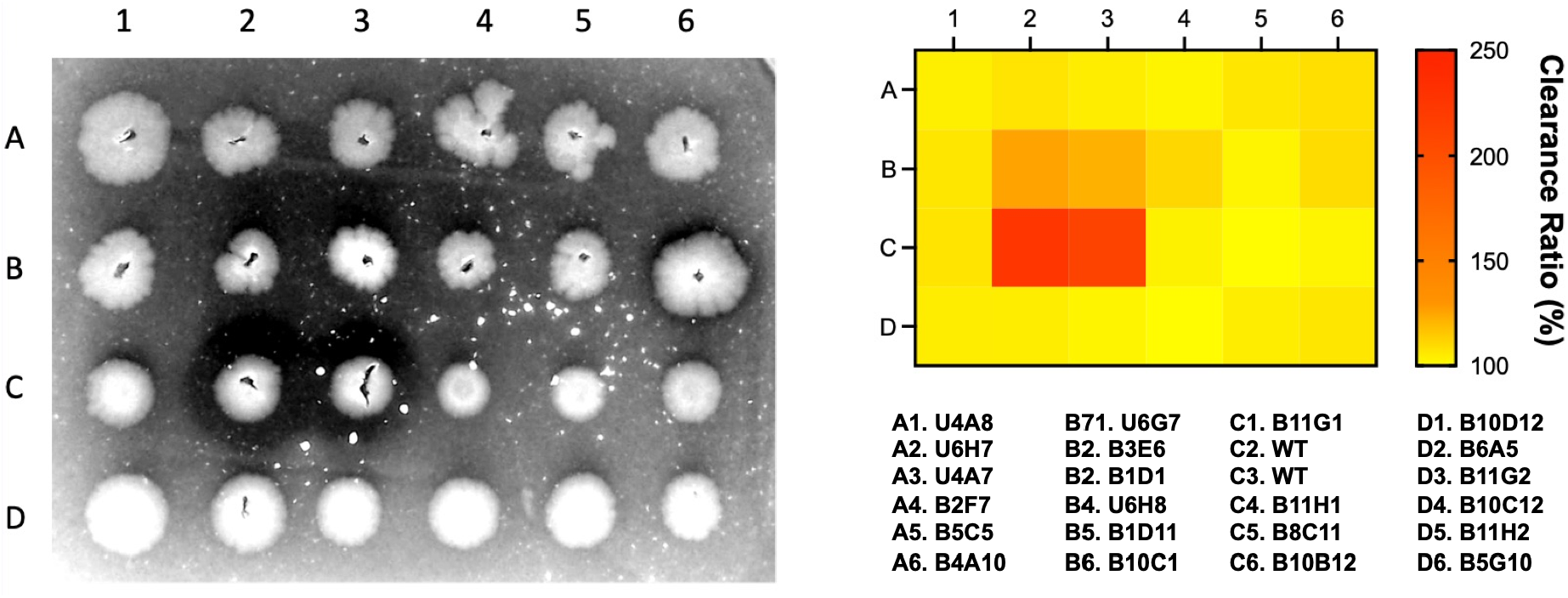
Rescreening of Selected PETase Mutants. Example data showing 23 mutants and WT FAST PETase with BHET clearance activity quantified using imaging algorithm.

Plasmids for the PETase mutants as well as WT FAST PETase and the TAG 82 negative control were transformed into *E. coli* BL21(DE3) which were grown in inducing media. The OD^600^ of the cultures were measured and their supernants were isolated and added to solutions of *p*-nitrophenyl butyrate (NO^2^-PhE) or 4-butoxyphenyl (2-methoxy-4-nitrophenyl)carbamate (NO^2^-PhU). Among the PETase mutants tested, Y82V had the most significant increase in normalized NO^2^-PhU hydrolysis activity (**Figure 6**).

**Figure 6.**
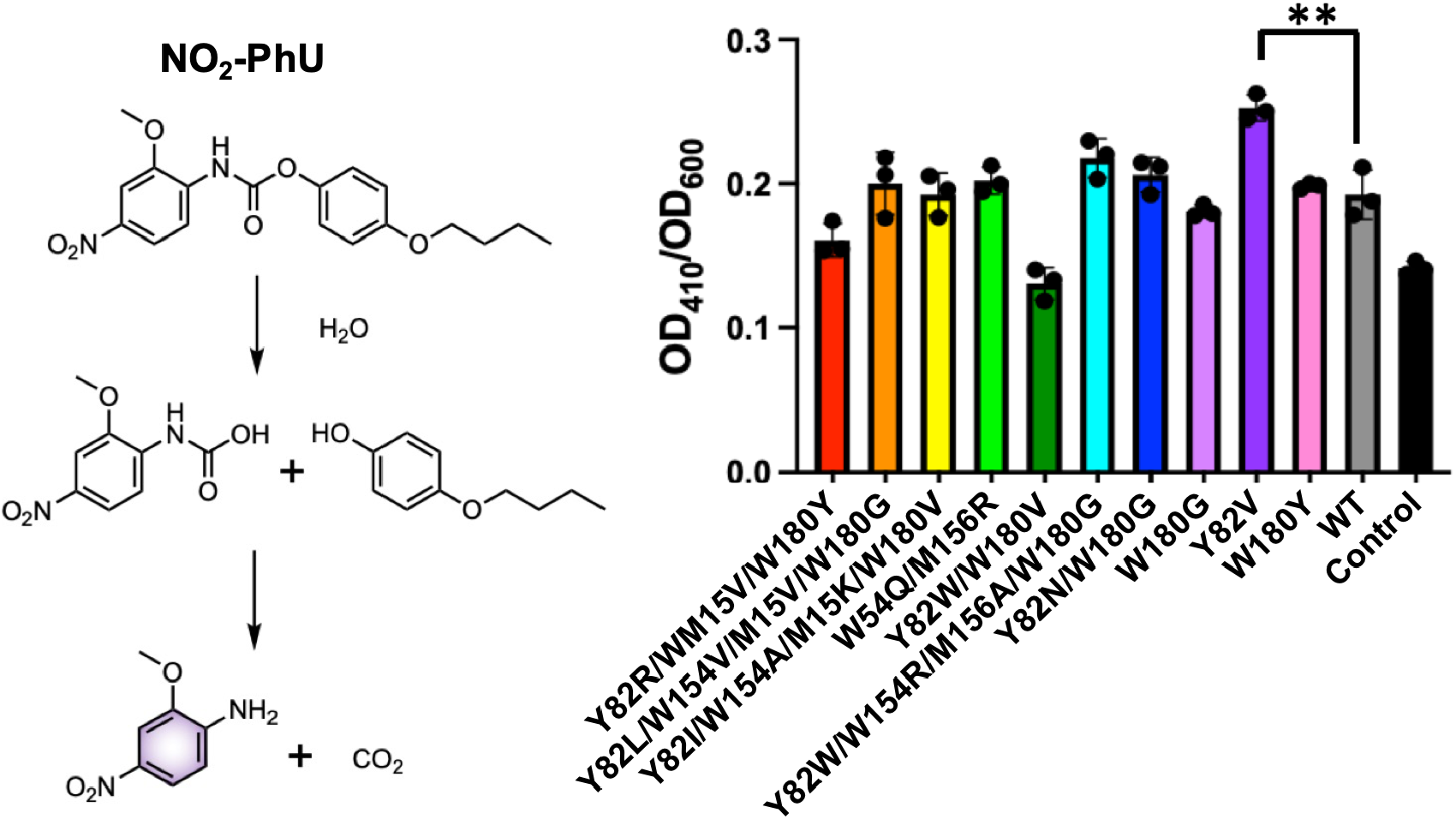
Urethane Cleavage Activity of PETase Mutants. OD^410^ measured 24 hours after mixing 100 µL of a saturated NO^2^-PhU in pH 8.0 Tris buffer with 100 µL concentrated supernatant from PETase mutant cultures. FAST and Y82TAG PETase cultures were used as controls. OD^410^ measurements were normalized by OD^600^ measurements of the cultures. ** = p<0.01

Retesting this mutant with supernatants adjusted for protein concentration showed that it had a 2.7-fold increase in NO^2^-PhU hydrolysis activity compared to the background-corrected activity of the parent WT FAST PETase (**Figure 7A**). Moreover, this mutant showed clearance activity on PU top-coated plates, where WT FAST PETase and the TAG82 control did not (**Figure 7B/C and** SI, **Figure S13**-**S14**).

**Figure 7.**
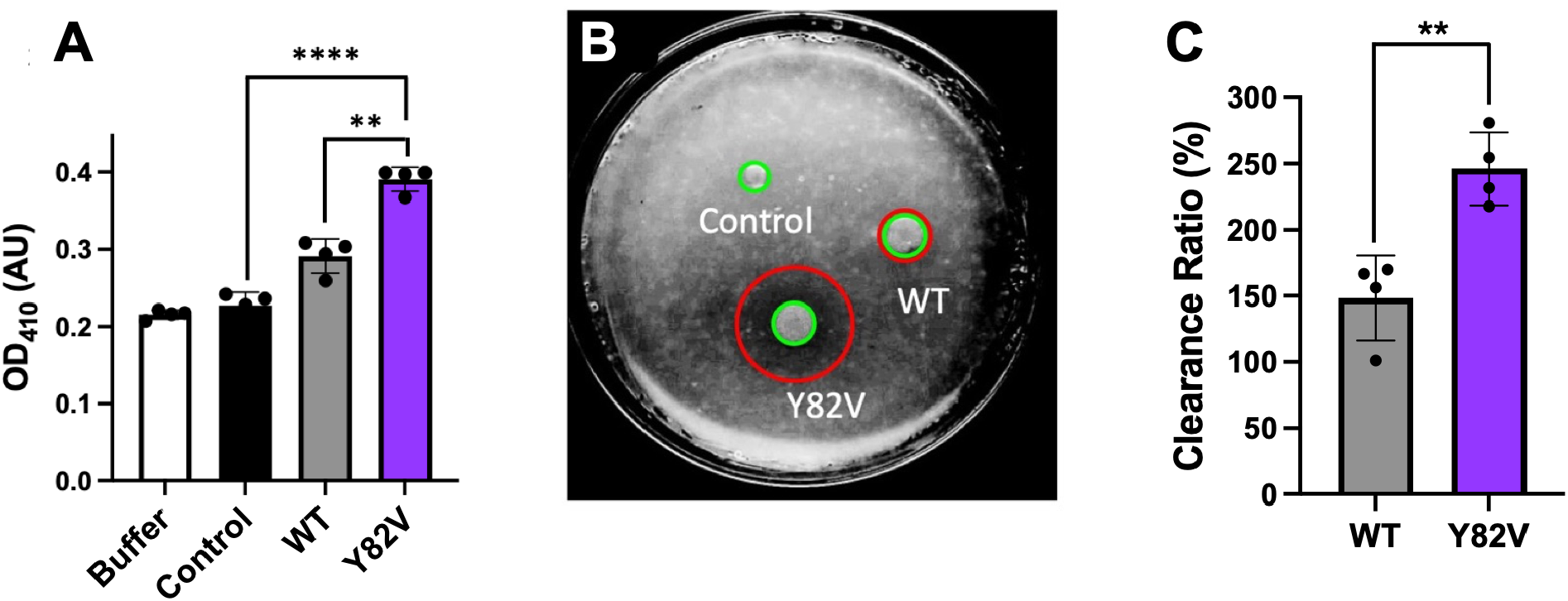
Urethane Cleavage Activity of Y82V PETase Mutant. **A.**OD^410^ measured 24 hours after mixing 100 µL of saturated NO^2^-PhU in pH 8.0 Tris buffer with 100 µL concentrated supernatant from PETase cultures. **B**. Image of colony growth and clearance on PU top-coated plate. **C**. PU clearance ratios for WT and Y82V mutant PETases. ** = p<0.01, **** = p<0.0001

To better understand the basis for the new urethanase activity of the Y82V PETase mutant, we generated computational models of the mutant protein with BHET or the NO^2^-PhU substrate bound based on the crystal structure of BHET bound to the parent FAST PETase (**Figure 8** and SI, **Figure S15**). Both substrates could be bound in the active site with the carbonyl bond oriented favorably for hydrolysis by the catalytic Ser residue. On the other hand, alignment of NO^2^-PhU in the FAST PETase active site identified steric clashes with M156 and Y82 that prevented favorable position of the carbamate for hydrolysis. This side of the active site can reorient to widen the active site slightly in the Y82V mutant due to the somewhat smaller sidechain of valine and repositioning of neighboring sidechains due to its β-branched structure.

**Figure 8.**
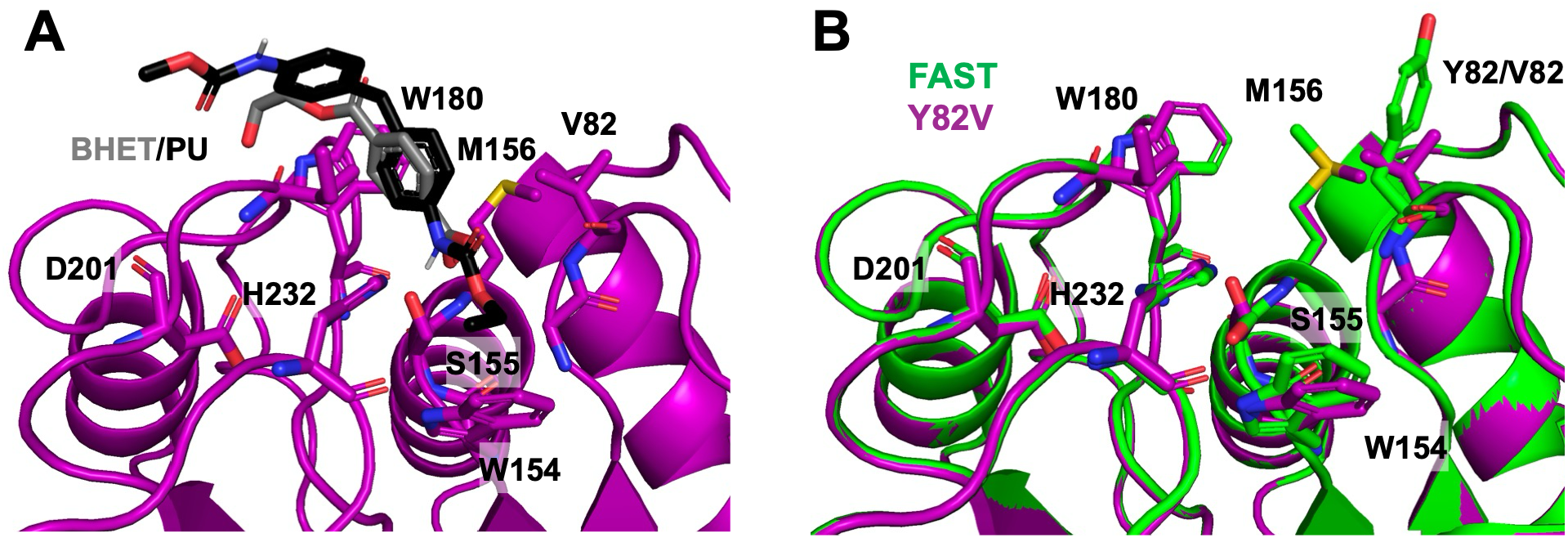
Structural Models of the Y82V FAST PETase Active Site. **A.**Model of the Y82V mutant after sculpting in PyMOL with PU substrate docked in the active site. BHET substrate from PDB ID 9LMV is shown to highlight relative substrate positioning. **B**. Overlay of Y82V mutant model and FAST PETase structure 9LMV highlighting movement of residues on the right side of the active site.

We also compared identified mutants in our screen to previously characterized PETase mutants and homologs. Wu et al analyzed the sequences of known PETases and used large language models to identify enzymes with PETase activity (Wu et al. 2025). The Y82V mutant and most of the mutants from our library with BHET clearance activity have amino acids that are not represented in their sequence database (**Figure 9** and SI, **Table S1**). For example, our selected sequences include Y82N and M154R mutations. This indicates that our mutations are accessing unusual parts of the PETase sequence space. Notably, there are putative PETases in their database with Thr, another beta-branched amino acid, at the equivalent of position 82.

**Figure 9.**
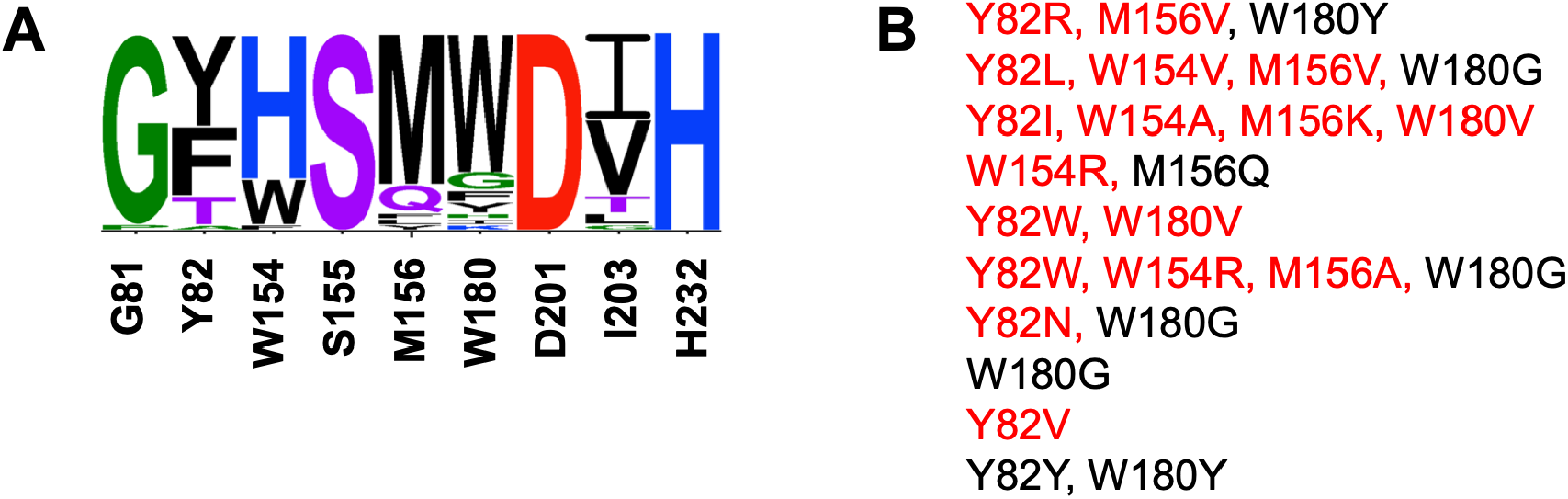
Sequence Analysis of Identified PETase Mutants. **A.**Sequence logo representation of 42 known PETases or related enzymes with putative PETase activity. Amino acid identities in the pelB-tagged FAST PETase parent construct are shown below. **B**. PETase variants from this screening assay with novel mutations highlighted in red.

## Conclusion

We successfully developed an enzyme that is effective at degrading both polyurethane and polyester plastic polymers. It shows significantly more activity than FAST-PETase, a more active version of the original Is PETase. Additionally, we successfully created an automated system that can be used to further increase activity through screening and selection. After optimization of the plate and growth conditions as well as the automated activity quantification algorithm, a 340,736-member mutant library was generated. Preliminary screening of >1000 colonies identified 23 mutants that showed activity on BHET or PU plates. These colonies were sequenced, and PE and PU degrading activity measured using chromophoric substrates, showing that one mutant, Y82V, acquired urethane degrading activity while maintaining ester degrading activity. The Y82V mutant was also tested on PU coated plates and showed the ability to break down actual polymeric PU. Although the increase in PU-degrading activity is modest, this is an exciting result from an initial small-scale screen. Implementation of robotic colony picking to more thoroughly explore sequence space in the 340,736-member mutant library may yield additional hits with greater PUase activity and nylonase activity as well.

The implications of this research are numerous. Expediated plastic degradation could not only significantly reduce pollution in landfills, but also in our habitats and other ecosystems. The need for incineration would also be significantly reduced, improving air quality. The bacteria can also be used to address microplastic issues in the sea and in food chains because they can detect and dispose of much smaller sized particles than sorting or filtering technology. Furthermore, the reduction of plastic waste accumulation would prevent spillover into wildlife areas and reduce the number of organisms that become entangled in microplastic spread.

## Acknowledgement

This research was supported by the University of Pennsylvania. C.J.P. thanks Riley Bednar for molecular biology advice and Andrew Zahrt for automation advice.

## Supporting Information

Additional figures, including sequence information, primary plate imaging data, additional structural and sequence analysis.

## Materials and Methods

### Materials and Instruments

Media components and antibiotics were purchased from Fisher Scientific (Hampton, NH, USA). Bis(2-Hydroxyethyl) terephthalate (BHET), nylon-6 pellets, *p*-nitrophenyl butyrate (NO^2^-PhE), and 4-butoxyphenyl (2-methoxy-4-nitrophenyl)carbamate (NO^2^-PhU) were purchased from Sigma-Aldrich (St. Loius, MO, USA). Low molecular weight polyurethane tent sealant solution was purchased from Gear Aid (Bellingham, WA, USA). Purified “Milli-Q” water used in media preparation was obtained from a Milli-Q 8 system (MilliporeSigma; Burlington, MA, USA). PCR reactions were performed with a BioRad T100 Thermocycler (Hercules, CA, USA). Electroporations were performed with a BioRad MicroPulser. BL21(DE3) *E. coli* cells were obtained from New England Biolabs (Ipswich, MA, USA) and electrocompetent preparations were made using published procedures (Gonzales et al. 2013). OmniTray™ single-well plates were purchased from Fisher Scientific. Greiner Bio-One clear bottom, black wall 96 well plates were purchased from Fisher Scientific. Optical density measurements were made using a Tecan Spark multimode platereader (Manendorf, Switzerland).

### Liquid Media

LB+Kan+IPTG liquid media was made with 10 g Tryptone, 10 g NaCl, and 5 g yeast extract, in 500 mL Milli-Q water (Luria Broth, LB). After autoclaving and cooling until warm to the touch, 500 µL kanamycin stock solution (Kan, 50 mg/mL in Milli-Q water) and 500 µL isopropyl β-D-1-thiogalactopyranoside stock solution (IPTG, 1 mM in Milli-Q water) were added to make LB+Kan+IPTG media.

### Solid Media

Bis(2-Hydroxyethyl) terephthalate (BHET) solid was ground into a fine powder using a mortar and pestle. Nylon was treated stirred with 18 M HCl at 120 °C in a round-bottomed flask until it formed a milky white solution. This was neutralized by dropwise addition of 1 M NaOH with pH monitored using pH strips. The neutralized solution was diluted to a final concentration of 0.2 g/mL using Milli-Q water. GearAid tent sealant solution was used as a source of low molecular weight polyurethane (PU).

LB+Kan+IPTG+BHET solid media was made with 10 g Tryptone, 10 g NaCl, 5 g yeast extract, and 15 g agar powder in 500 mL Milli-Q water. After autoclaving and cooling until warm to the touch, 500 µL Kan stock solution and 500 µL IPTG sock solution were added to make LB+Kan+IPTG media and then portioned into smaller flasks. Ground BHET powder was added to each flask at various final concentrations and the flask swirled to suspend it uniformly. Petri dishes were then filled to a 0.5 cm depth using a serological pipet and stored at 4 °C after solidifying.

LB+Kan+IPTG+Nylon solid media was made similarly to the BHET media, with the neutralized nylon solution added to LB+Kan+IPTG media to a 10% or 20% (v/v) final concentration and the flask swirled to suspend it uniformly. Petri dishes were then filled to a 0.5 cm depth and stored at 4 °C after solidifying.

LB+Kan+IPTG+PU solid media was made by filling Petri dishes to 0.5 cm depth with LB+Kan+IPTG media, allowing it to solidify, and then coating the surface with a 1%, 2%, or 5% dilutions of PU coating solution in LB+Kan+IPTG liquid media. The Petri dishes were stored at 4 °C after the PU topcoat solidified.

### Plasmid Design

Obtained Is, Hot, and FAST PETase sequences from Carter et al. (Carter et al. 2024), Thermo PETase sequence from Son et al. (Son et al. 2019), Dura PETase sequence from Liu et al. (Liu et al. 2022), and NylB sequence from Bell *et al*. (Bell et al. 2024), designed G-blocks for insertion between XbaI and XhoI sites in pET28a(+) in SnapGene and ordered plasmids from Genscript, with sequences confirmed by whole plasmid sequencing.

### Y82TAG Negative Control Plasmid

A TAG mutation was introduced using the primers below to introduce a stop codon so that the plasmid could serve as an inactive background for library generation and plate-based screening.

**Y82-TAG For:** TTGTCCCCGGCTAGACTGCTCGCCA

**Y82-TAG Rev:** TGGCGAGCAGTCTAGCCGGGGACAA

### Four Position Mutant Library Construction

Primers were designed for saturation mutagenesis at W82, W154, M156, and W180, using primer with “22 codon trick” mixtures (Kille et al. 2013) for positions 82, 154, and 180 and a standard NNK mixture at position 156. The primer sequences given below were purchased from IDT and dissolved in Milli-Q water to a final concentration of 200 ng/mL in a 12:9:1 ratio of the NDT, VHG, and TGG forward primers. The reverse primers were dissolved in Milli-Q water to a final concentration of 200 ng/mL. The DNA insert was prepared in a series of PCR reactions as shown below. First, four regions of the insert were generated by PCR amplification of the pET28a-FAST plasmid region between the For and Rev sequences shown below. Blocks 1, 2, 3, and 4 were gel purified and used in further rounds of PCR amplification. A combined 1+2 block was amplified using the XbaI For and W154M156 Rev primers. A combined 3+4block was amplified using the W154M156 For and XhoI Rev primers. The sequence of the primers were:

**Y82-NDT Rev:** CCACCATTTGATACTGCTTTGGCGAGCAGTAHNGCCGGGGACAA **Y82-VHG Rev:** CCACCATTTGATACTGCTTTGGCGAGCAGTCDBGCCGGGGACAA **Y82-TGG Rev:** CCACCATTTGATACTGCTTTGGCGAGCAGTCCAGCCGGGGACAA

**Y82-Lib For:** ACTGCTCGCCAAAGCAGTATCAAATGGTGG

**W154M156-NDT Rev:** GGAAATAAGGCTCCCCCCACCGCCAHNGCTMNNTCCCATAA **W154M156-VHG Rev:** GGAAATAAGGCTCCCCCCACCGCCCDBGCTMNNTCCCATAA **W154M156-TGG Rev:** GGAAATAAGGCTCCCCCCACCGCCCCAGCTMNNTCCCATAA

**W154M156-Lib For:** GGCGGTGGGGGGAGCCTTATTTCC

**W180-NDT Rev:** GGACGGTGACCGAGCTGAAATTAGTTGACGAATGAHNCGGGGCCTGAGG **W180-VHG Rev:** GGACGGTGACCGAGCTGAAATTAGTTGACGAATGCDBCGGGGCCTGAGG **W180-TGG Rev:** GGACGGTGACCGAGCTGAAATTAGTTGACGAATGCCACGGGGCCTGAGG

**W180-Lib For:** CATTCGTCAACTAATTTCAGCTCGGTCACCGTCC

**XbaI For:** AATTGTGAGCGGATAACAATTCCCC

**XhoI Rev:** GGGCTTTGTTAGCAGCCGGATCTCA

### Library Growth and Screening Procedure

Gibson assembly reactions were transformed into electrocompetent BL21(DE3) *E. coli* cells using a BioRad electroporator. After recovery for 60-90 min in 500 µL SOC media, they were concentrated to ∼50 µL by centrifugation, removal of the supernatant, and resuspension, then they were plated on LB+Kan plates for overnight growth. On the following day, colonies were picked and grown in 1 mL LB+Kan liquid media in a 96 well plate (2 mL wells) with 250 rpm shaking at 37 °C. After overnight growth, a 48 pin replicator was used to plate colonies on LB+Kan+IPTG+BHET and LB+Kan+IPTG+PU plates. The remaining cultures were mixed with 50% glycerol (v/v in Milli-Q water) and stored as 5 cryo-preserved stocks in 96 well plates at -80°C.

LB+Kan+IPTG+BHET and LB+Kan+IPTG+PU plates were incubated overnight at 30 °C or 37 °C and imaged after 1, 2, and 3 days. BHET or PU clearance activity was quantified using the image analysis algorithm. All colonies showing BHET activity above our threshold were selected for sequencing and further analysis.

### Plate Imaging

The custom imaging setup was constructed with a light pad (Portable A4 LED copy board, LitEnergy; Wewahitchka, FL, USA) placed flat on the benchtop set to maximum brightness with a 10 x 15 cm black paper rectangle placed in the center and a 3D printed rack with a slot for rectangular media plates, elevating the 6 cm above the paper and allowing light to pass through. The camera (iPhone 12, Apple Inc.; Cupertino, CA, USA) was mounted to a ring stand 20 cm above the plate. Room lights were turned off to minimize glare.

### Classical Image Analysis

Classical image analysis was used to quantify bacterial colony growth on solid culture plates. Plate images were acquired under consistent lighting and camera distance to reduce variation between samples. Images were imported into Python and preprocessed by converting to grayscale, adjusting contrast, and reducing noise with a gaussian blur. The plate region was identified using a Hough circles detection to exclude background outside the agar surface. CLAHE (Contrast Limited Adaptive Histogram Equalization) was used to make boundaries easier to distinguish. Brightness thresholding was used to identify colonies using Open CV’s

SimpleBlobDetector. Then radius was calculated using the equation 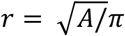 where A is the bright area in pixels. Finally, radial intensity profiling was used to identify halo clearance areas. Starting at the center of the predicted colony, pixel brightness was mapped. Those with a decrease after the approximated radius of the colony and followed with a less intense increase after a number of pixels were identified as having a halo. After 5 trials were run, each at different angles from the radius, the halo radius was averaged. The ratio of the colony radius to the halo radius was used to quantify activity. The total number of colonies and distribution of colony sizes were calculated for each image and used to compare bacterial growth across conditions.

### Colony Sequencing

Selected colonies were picked, grown in 500 µL LB+Kan liquid media with 250 rpm shaking for 6-8 h to OD^600^ ∼0.5, and 100 µL of the turbid media was submitted for whole plasmid sequencing. The remaining media was mixed with 400 µL of 50% glycerol (v/v in Milli-Q water) and frozen at -80 °C. If sequencing errors were observed, the colony was grown in a 5 mL LB+Kan culture and a miniprep procedure performed to isolate plasmid DNA for whole plasmid sequencing.

Whole plasmid sequencing was used to confirm PETase gene insertion and identify mutations. The resulting sequence chromatograms were reviewed for read quality. The presence, orientation, and full or partial integrity of the insert were assessed by comparison to the starting PETase construct. Base substitutions, insertions, deletions, or other sequence deviations were recorded. Constructs that matched the expected sequence or contained only acceptable intended changes were classified as successful, while constructs with frameshifts, incorrect insertions, or unintended mutations were excluded from further analysis.

### Solution-Phase Esterase and Urethanase Activity Screening

Glycerol stocks for colonies selected from BHET and PU plate screening were used to inoculate 5 mL cultures of LB+Kan+IPTG liquid media. These cultures were grown for 8 h and the OD^600^ measured for 100 µL aliquots on a Tecan multimode platereader. Then the cells were pelleted and the supernatant was collected and concentrated to 500 µL using an Amicon 3kD MWCO spin filter. To measure esterase activity, 100 µL of the concentrated supernatant was mixed with 100 µL of a saturated solution of *p*-nitrophenyl butyrate (NO^2^-PhE) in pH 8.0 Tris buffer. FAST and Y82TAG PETase cultures were used as controls. The absorbance at 415 nm was measured with 250 rpm shaking. To measure urethanase activity, 100 µL of the concentrated supernatant was mixed with 100 µL of a saturated solution of 4-butoxyphenyl (2-methoxy-4-nitrophenyl)carbamate (NO^2^-PhU) in pH 8.0 Tris buffer. FAST and Y82TAG PETase cultures were used as controls. The absorbance at 410 nm was measured with 250 rpm shaking.

### One way ANOVA and student’s t-tests to determine significance of differences

Statistical analysis was performed in Prism 9 (GraphPad Software, San Diego, CA, USA) to evaluate whether measured differences among enzymes or mutants were significant. For comparisons involving more than two groups, a one-way ANOVA was conducted using replicate measurements for each enzyme. Before analysis, data were checked for obvious outliers and summarized as mean ± standard deviation. If the ANOVA indicated a significant overall difference among groups, pairwise comparisons were performed using Student’s t-tests to identify which enzymes differed from one another. When appropriate, multiple-comparison correction was applied to reduce false-positive results. For direct comparison between two groups only, a two-tailed Student’s t-test was used. A significance threshold of p < 0.05 was used unless otherwise specified.

### Computational Modeling

Atomic coordinates for a FAST PETase mutant under PDB ID 9LMV (Li et al. 2025) were downloaded from the Protein Data Bank (PDB), imported into PyMOL (Schrödinger Inc.; New York, NY, USA) and Y82V mutation was made using the standard protein mutagenesis function. Polyurethane (PU) monomer unit generated in Chemdraw (Revvity Signals Software; Waltham, MA, USA), exported as SMILES string and 3D conformer generated by NovoPro (https://www.novoprolabs.com/tools/smiles2pdb) as a pdb file.

PU SMILES: O=C(OCCO)NC(C=C1)=CC=C1CC2=CC=C(NC(OCCCO)=O)C=C2

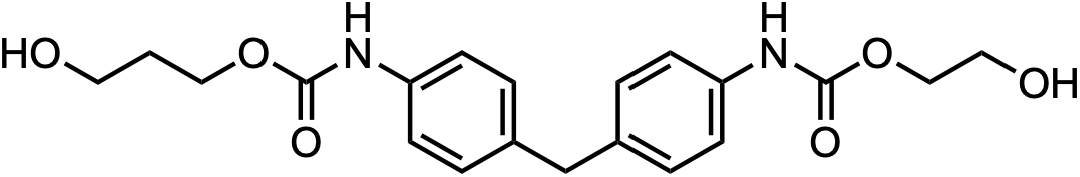

PU pdb file imported into Y82V structure in PyMOL, manually aligned to BHET structure, and local minimizations were performed with the Sculpting tool.

### Sequence Analysis

Sequences were obtained from Wu et al. (Wu et al. 2025), aligned, and used to generate logo IDs with WebLogo (https://weblogo.threeplusone.com) (Crooks et al. 2004).

