## Supporting Information for "Directed Evolution of Multimodal Plastic Degrading Enzymes"

**Table of Contents**

PETase and nylonase plasmid design …………………………………………………………… S2

Images from initial testing of BHET, nylon, and PU plates …………………………………..… S6

Image analysis algorithm ……………………………………………………….…………….…… S7

Images of library plates ……………………………………………………….…………………… S8

Sequences of selected clones ………………………………………….…….………………… S11

PUase Activity Analysis ……………………………………..…….…………..…………… S15

Computational Models ……………………..……………………..…….………….…………… S16

PETase Sequences ……………………………..……………..…….…………..…………… S17

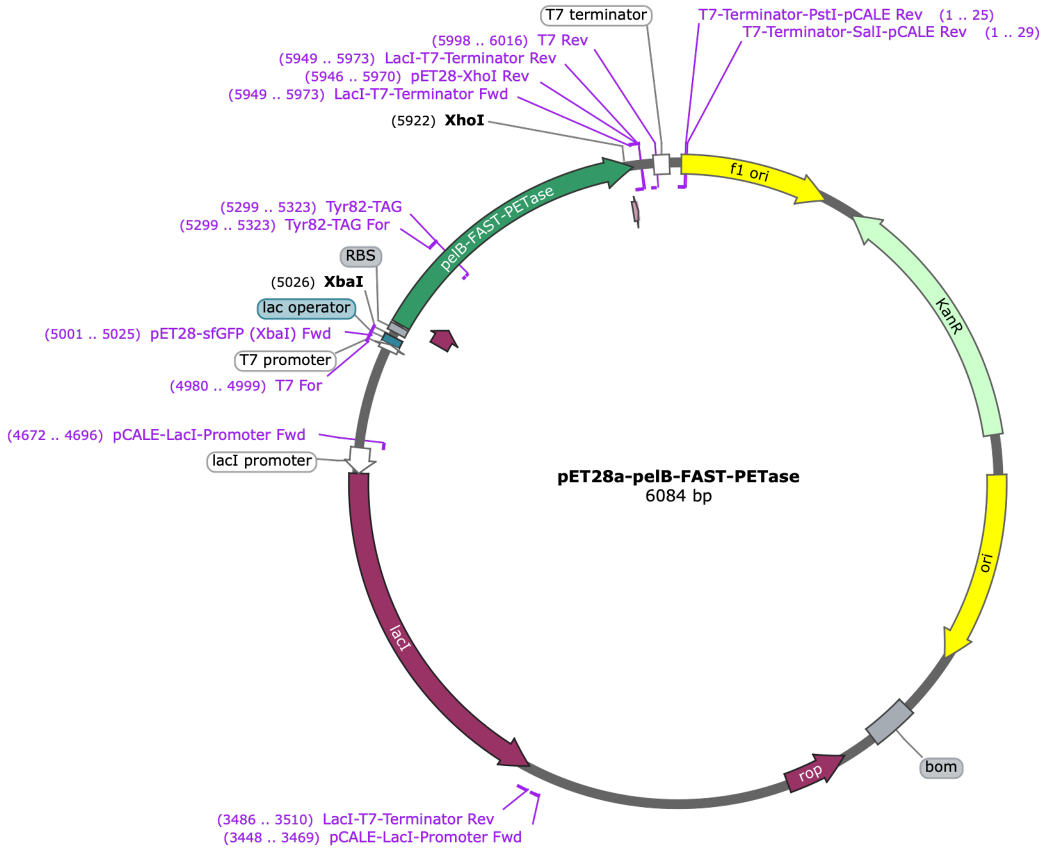


**Figure S1.** General Plasmid Map.

**pelB-FAST-PETase G-block (XbaI for and XhoI rev primer sequences underlined):**
<u>AATTGTGAGCGGATAACAATTCCCC</u>TCTAGAAATAATTTTGTTTAACTTTAAGAAGGAGATATACCATG
AAATACCTGCTGCCGACCGCTGCTGCTGGTCTGCTGCTTCTCGCTGCCCAGCCGGCGATGGCCCA
GACTAATCCATACGCCCGCGGACCTAACCCTACGGCGGCGAGTTTGGAGGCTTCTGCTGGTCCGTT
CACAGTTCGTTCTTTTACGGTGTCGCGTCCAAGCGGGTACGGCGCTGGAACCGTCTACTATCCCAC
TAACGCTGGAGGCACAGTAGGAGCTATCGCCATTGTCCCCGGCTACACTGCTCGCCAAAGCAGTAT
CAAATGGTGGGGTCCGCGCTTGGCCTCACATGGGTTTGTTGTGATTACCATCGACACTAATTCCACA
TTGGACCAGCCAGAATCACGTTCTTCACAACAAATGGCTGCCTTACGCCAAGTGGCGTCGCTTAATG
GGACTTCTAGTTCACCAATCTACGGTAAGGTAGATACTGCTCGCATGGGAGTTATGGGATGGAGCAT
GGGCGGTGGGGGGAGCCTTATTTCCGCAGCGAATAACCCATCCCTGAAGGCTGCCGCACCTCAGG
CCCCGTGGCATTCGTCAACTAATTTCAGCTCGGTCACCGTCCCTACTCTTATTTTCGCTTGTGAAAAC
GACAGTATCGCACCTGTTAATTCATCGGCACTGCCAATCTATGATTCCATGTCCCAGAACGCTAAGC
AATTCCTGGAGATTAAAGGGGGAAGTCACTCGTGCGCTAATAGTGGGAACAGTAATCAAGCTCTTAT
CGGGAAGAAGGGTGTTGCATGGATGAAACGCTTTATGGACAACGACACACGTTATAGTACCTTTGCG
TGCGAAAATCCTAATTCTACGGCGGTGAGTGACTTCCGCACGGCAAATTGCAGCCTCGAGCACCAC
CACCACCACCAC<u>TGAGATCCGGCTGCTAACAAAGCCC</u>

**pelB-FAST-PETase Protein Sequence (pelB underlined):**
<u>MKYLLPTAAAGLLLLAAQPAMA</u>QTNPYARGPNPTAASLEASAGPFTVRSFTVSRPSGYGAGTVYYPTNA
GGTVGAIAIVPGYTARQSSIKWWGPRLASHGFVVITIDTNSTLDQPSSRSSQQMAALRQVASLNGTSSSP
IYGKVDTARMGVMGWSMGGGGSLISAANNPSLKAAAPQAPWDSSTNFSSVTVPTLIFACENDSIAPVNS
SALPIYDSMSRNAKQFLEINGGSHSCANSGNSNQALIGKKGVAWMKRFMDNDTRYSTFACENPNSTRV
SDFRTANCSLEHHHHHH

**Figure S2.** FAST PETase DNA and Protein Sequences.

**pelB-Is-PETase G-block (XbaI for and XhoI rev primer sequences underlined:**
<u>AATTGTGAGCGGATAACAATTCCCC</u>TCTAGAAATAATTTTGTTTAACTTTAAGAAGGAGATATACCATG
AAATACCTGCTGCCGACCGCTGCTGCTGGTCTGCTGCTTCTCGCTGCCCAGCCGGCGATGGCCCA
GACTAATCCATACGCCCGCGGACCTAACCCTACGGCGGCGAGTTTGGAGGCTTCTGCTGGTCCGTT
CACAGTTCGTTCTTTTACGGTGTCGCGTCCAAGCGGGTACGGCGCTGGAACCGTCTACTATCCCAC
TAACGCTGGAGGCACAGTAGGAGCTATCGCCATTGTCCCCGGCTACACTGCTCGCCAAAGCAGTAT
CAAATGGTGGGGTCCGCGCTTGGCCTCACATGGGTTTGTTGTGATTACCATCGACACTAATTCCACA
TTGGACCAGCCAAGCTCACGTTCTTCACAACAAATGGCTGCCTTACGCCAAGTGGCGTCGCTTAATG
GGACTTCTAGTTCACCAATCTACGGTAAGGTAGATACTGCTCGCATGGGAGTTATGGGATGGAGCAT
GGGCGGTGGGGGGAGCCTTATTTCCGCAGCGAATAACCCATCCCTGAAGGCTGCCGCACCTCAGG
CCCCGTGGGATTCGTCAACTAATTTCAGCTCGGTCACCGTCCCTACTCTTATTTTCGCTTGTGAAAAC
GACAGTATCGCACCTGTTAATTCATCGGCACTGCCAATCTATGATTCCATGTCCCGCAACGCTAAGC
AATTCCTGGAGATTAACGGGGGAAGTCACTCGTGCGCTAATAGTGGGAACAGTAATCAAGCTCTTAT
CGGGAAGAAGGGTGTTGCATGGATGAAACGCTTTATGGACAACGACACACGTTATAGTACCTTTGCG
TGCGAAAATCCTAATTCTACGCGCGTGAGTGACTTCCGCACGGCAAATTGCAGCCTCGAGCACCAC
CACCACCACCAC<u>TGAGATCCGGCTGCTAACAAAGCCC</u>

**pelB-Is-PETase Protein Sequence (pelB underlined):**
<u>MKYLLPTAAAGLLLLAAQPAMA</u>QTNPYARGPNPTAASLEASAGPFTVRSFTVSRPSGYGAGTVYYPTNA
GGTVGAIAIVPGYTARQSSIKWWGPRLASHGFVVITIDTNSTLDQPSSRSSQQMAALRQVASLNGTSSSP
IYGKVDTARMGVMGWSMGGGGSLISAANNPSLKAAAPQAPWDSSTNFSSVTVPTLIFACENDSIAPVNS
SALPIYDSMSRNAKQFLEINGGSHSCANSGNSNQALIGKKGVAWMKRFMDNDTRYSTFACENPNSTRV
SDFRTANCSLEHHHHHH

**Figure S3.** Is PETase DNA and Protein Sequences.

**pelB-Dura-PETase G-block (XbaI for and XhoI rev primer sequences underlined:**
AATTGTGAGCGGATAACAATTCCCCTCTAGAAATAATTTTGTTTAACTTTAAGAAGGAGATATACCATG
AAATACCTGCTGCCGACCGCTGCTGCTGGTCTGCTGCTTCTCGCTGCCCAGCCGGCGATGGCCCA
GACTAATCCATACGCCCGCGGACCTAACCCTACGGCGGCGAGTTTGGAGGCTTCTGCTGGTCCGTT
CACAGTTCGTTCTTTTACGGTGTCGCGTCCAAGCGGGTACGGCGCTGGAACCGTCTACTATCCCAC
TAACGCTGGAGGCACAGTAGGAGCTATCGCCATTGTCCCCGGCTACACTGCTCGCCAAAGCAGTAT
CAAATGGTGGGGTCCGCGCTTGGCCTCACATGGGTTTGTTGTGATTACCATCGACACTAATTCCACA
TTTGACTATCCAAGCTCACGTTCTTCACAACAAATGGCTGCCTTACGCCAAGTGGCGTCGCTTAATG
GGGATTCTAGTTCACCAATCTACGGTAAGGTAGATACTGCTCGCATGGGAGTTATGGGACATAGCAT
GGGCGGTGGGGCGAGCCTTCGCTCCGCAGCGAATAACCCATCCCTGAAGGCTGCCATTCCTCAGG
CCCCGTGGGATTCGCAGACTAATTTCAGCTCGGTCACCGTCCCTACTCTTATTTTCGCTTGTGAAAA
CGACAGTATCGCACCTGTTAATTCACATGCACTGCCAATCTATGATTCCATGTCCCGCAACGCTAAG
CAATTCCTGGAGATTAACGGGGGAAGTCACTCGTGCGCTAATAGTGGGAACAGTAATCAAGCTCTTA
TCGGGAAGAAGGGTGTTGCATGGATGAAACGCTTTATGGACAACGACACACGTTATAGTACCTTTGC
GTGCGAAAATCCTAATTCTACGGCGGTGAGTGACTTCCGCACGGCAAATTGCAGCCTCGAGCACCA
CCACCACCACCAC<u>TGAGATCCGGCTGCTAACAAAGCCC</u>

**pelB-Dura-PETase Protein Sequence (pelB underlined):**
<u>MKYLLPTAAAGLLLLAAQPAMA</u>QTNPYARGPNPTAASLEASAGPFTVRSFTVSRPSGYGAGTVYYPTNA
GGTVGAIAIVPGYTARQSSIKWWGPRLASHGFVVITIDTNSTFDYPSSRSSQQMAALRQVASLNGDSSSP
IYGKVDTARMGVMGHSMGGGASLRSAANNPSLKAAIPQAPWDSQTNFSSVTVPTLIFACENDSIAPVNS
HALPIYDSMSRNAKQFLEINGGSHSCANSGNSNQALIGKKGVAWMKRFMDNDTRYSTFACENPNSTAV
SDFRTANCSLEHHHHHH

**Figure S4.** Dura PETase DNA and Protein Sequences.

**pelB-Thermo-PETase G-block (XbaI for and XhoI rev primer sequences underlined:**
AATTGTGAGCGGATAACAATTCCCCTCTAGAAATAATTTTGTTTAACTTTAAGAAGGAGATATACCATG
AAATACCTGCTGCCGACCGCTGCTGCTGGTCTGCTGCTTCTCGCTGCCCAGCCGGCGATGGCCCA
GACTAATCCATACGCCCGCGGACCTAACCCTACGGCGGCGAGTTTGGAGGCTTCTGCTGGTCCGTT
CACAGTTCGTTCTTTTACGGTGTCGCGTCCAAGCGGGTACGGCGCTGGAACCGTCTACTATCCCAC
TAACGCTGGAGGCACAGTAGGAGCTATCGCCATTGTCCCCGGCTACACTGCTCGCCAAAGCAGTAT
CAAATGGTGGGGTCCGCGCTTGGCCTCACATGGGTTTGTTGTGATTACCATCGACACTAATTCCACA
TTGGACCAGCCAGAATCACGTTCTTCACAACAAATGGCTGCCTTACGCCAAGTGGCGTCGCTTAATG
GGACTTCTAGTTCACCAATCTACGGTAAGGTAGATACTGCTCGCATGGGAGTTATGGGATGGAGCAT
GGGCGGTGGGGGGAGCCTTATTTCCGCAGCGAATAACCCATCCCTGAAGGCTGCCGCACCTCAGG
CCCCGTGGCATTCGTCAACTAATTTCAGCTCGGTCACCGTCCCTACTCTTATTTTCGCTTGTGAAAAC
GACAGTATCGCACCTGTTAATTCATCGGCACTGCCAATCTATGATTCCATGTCCCGCAACGCTAAGC
AATTCCTGGAGATTAACGGGGGAAGTCACTCGTGCGCTAATAGTGGGAACAGTAATCAAGCTCTTAT
CGGGAAGAAGGGTGTTGCATGGATGAAACGCTTTATGGACAACGACACACGTTATAGTACCTTTGCG
TGCGAAAATCCTAATTCTACGGCGGTGAGTGACTTCCGCACGGCAAATTGCAGCCTCGAGCACCAC
CACCACCACCACTGAGATCCGGCTGCTAACAAAGCCC

**pelB-Thermo-PETase Protein Sequence (pelB underlined):**
MKYLLPTAAAGLLLLAAQPAMAQTNPYARGPNPTAASLEASAGPFTVRSFTVSRPSGYGAGTVYYPTNA
GGTVGAIAIVPGYTARQSSIKWWGPRLASHGFVVITIDTNSTLDQPESRSSQQMAALRQVASLNGTSSSP
IYGKVDTARMGVMGWSMGGGGSLISAANNPSLKAAAPQAPWHSSTNFSSVTVPTLIFACENDSIAPVNS
SALPIYDSMSRNAKQFLEINGGSHSCANSGNSNQALIGKKGVAWMKRFMDNDTRYSTFACENPNSTAVS
DFRTANCSLEHHHHHH

**Figure S5.** Thermo PETase DNA and Protein Sequences

**pelB-Hot-PETase G-block (XbaI for and XhoI rev primer sequences underlined:**
AATTGTGAGCGGATAACAATTCCCCTCTAGAAATAATTTTGTTTAACTTTAAGAAGGAGATATACCATG
AAATACCTGCTGCCGACCGCTGCTGCTGGTCTGCTGCTTCTCGCTGCCCAGCCGGCGATGGCCCA
GACTAATCCATACGCCCGCGGACCTAACCCTACGGCGGCGAGTTTGGAGGCTTCTGCTGGTCCGTT
CACAGTTCGTTCTTTTACGGTGGCGCGTCCAGTGGGGTACGGCGCTGGAACCGTCTACTATCCCAC
TAACGCTGGAGGCACAGTAGGAGCTATCGCCATTGTCCCCGGCTACACTGCTACCCAAAGCAGTAT
CAACTGGTGGGGTCCGCGCTTGGCCTCACATGGGTTTGTTGTGATTACCATCGACACTAATTCCACA
TTGGACAAACCAGAATCACGTTCTTCACAACAAATGGCTGCCTTACGCCAAGTGGCGTCGCTTAATG
GGACTTCTAGTTCACCAATCTACGGTAAGGTAGATACTGCTCGCGGCGGAGTTATGGGATGGAGCA
TGGGCGGTGGGGGGAGCCTTATTTCCGCAGCGAATAACCCATCCCTGAAGGCTGCCGCAGTGATG
GCCCCGTGGCATTCGTCAACTAATTTCAGCTCGGTCACCGTCCCTACTCTTATTTTCGCTTGTGAAAA
CGACCGCATCGCACCTGTTAAAGAATATGCACTGCCAATCTATGATTCCATGTCCCTGAACGCTAAG
CAATTCCTGGAGATTTGCGGGGGAAGTCACTCGTGCGCTTGCAGTGGGAACAGTAATCAAGCTCTT
ATCGGGATGAAGGGTGTTGCATGGATGAAACGCTTTATGGACAACGACACACGTTATAGTCAGTTTG
CGTGCGAAAATCCTAATTCTACGGCGGTGTGCGACTTCCGCACGGCAAATTGCAGCCTCGAGCACC
ACCACCACCACCACTGAGATCCGGCTGCTAACAAAGCCC

**pelB-Hot-PETase Protein Sequence (pelB underlined):**
MKYLLPTAAAGLLLLAAQPAMAQTNPYARGPNPTAASLEASAGPFTVRSFTVARPVGYGAGTVYYPTNA
GGTVGAIAIVPGYTATQSSINWWGPRLASHGFVVITIDTNSTLDKPESRSSQQMAALRQVASLNGTSSSPI
YGKVDTARGGVMGWSMGGGGSLISAANNPSLKAAAVMAPWHSSTNFSSVTVPTLIFACENDRIAPVKEY
ALPIYDSMSLNAKQFLEICGGSHSCACSGNSNQALIGMKGVAWMKRFMDNDTRYSQFACENPNSTAVC
DFRTANCSLEHHHHHH

**Figure S6.** Hot PETase DNA and Protein Sequences

**pelB-NylB G-block (XbaI for and XhoI rev primer sequences underlined:**
<u>AATTGTGAGCGGATAACAATTCCCC</u>TCTAGAAATAATTTTGTTTAACTTTAAGAAGGAGATATACCATG
AAATACCTGCTGCCGACCGCTGCTGCTGGTCTGCTGCTTCTCGCTGCCCAGCCGGCGATGGCCATG
AACGCTCGGTCGACGGGACAGCATCCGGCGCGTTATCCGGGCGCGGCCGCAGGAGAACCTACTCT
GGATTCGTGGCAGGAAGCCCCGCACAACCGCTGGGCCTTTGCGCGTCTGGGTGAGCTGCTTCCCA
CAGCGGCTGTGTCAAGACGTGATCCGGCGACACCAGCGGAACCGGTTGTTCGGCTGGACGCGTTA
GCGACACGCCTGCCGGATCTGGAACAGCGCTTAGAGGAAACCTGCACCGACGCATTTCTCGTTTTA
AGAGGGTCAGAAGTATTAGCTGAATATTATCGTGCAGGGTTTGCTCCGGATGACCGTCATCTTCTTA
TGTCTGTGAGTAAATCCTTATGCGGCACTGTCGTGGGCGCTCTGATTGACGAAGGCAGAATAGATCC
GGCTCAGCCCGTGACTGAGTATGTCCCAGAACTTGCCGGGAGTGTCTATGATGGCCCGAGCGTGTT
ACAGGTCCTGGATATGCAGATATCTATCGATTATAACGAGGATTATGTGGATCCTGCTAGCGAAGTG
CAGACACATGACCGTTCAGCAGGTTGGCGCACCCGTCGCGATGGAGATCCAGCCGATACATATGAA
TTCCTGACTACCCTGCGTGGCGATGGTGGTACGGGAGAGTTTCAATATTGTTCGGCGAACACAGAT
GTTCTCGCGTGGATAGTCGAACGTGTAACTGGACTGCGCTATGTTGAGGCGCTTAGCACATATCTCT
GGGCTAAGTTGGACGCGGATCGCGATGCTACTATTACGGTGGATCAAACGGGTTTTGGATTTGCAA
ATGGTGGCGTGAGTTGTACAGCGCGGGATCTGGCACGGGTAGGCCGGATGATGCTGGATGGAGGG
GTTGCCCCGGGGGGCCGTGTAGTAAGTCAGGGGTGGGTGGAATCCGTCCTTGCGGGGGGTTCGC
GGGAGGCTATGACTGACGAAGGCTTTACCTCAGCGTTTCCCGAAGGTAGCTACACGCGACAGTGGT
GGTGTACTGGGAACGAACGGGGTAACGTATCTGGAATTGGTATCCATGGACAGAACTTATGGCTGG
ATCCTCGTACCGACTCCGTGATAGTTAAACTTTCATCATGGCCAGATCCCGATACACGCCATTGGCA
CGGCTTGCAATCAGGGATTTTACTGGACGTTTCCCGTGCTCTTGATGCTGTACTCGAGCACCACCAC
CACCACCAC<u>TGAGATCCGGCTGCTAACAAAGCCC</u>

**pelB-NylB Protein Sequence (pelB underlined):**
<u>MKYLLPTAAAGLLLLAAQPAMA</u>MNARSTGQHPARYPGAAAGEPTLDSWQEAPHNRWAFARLGELLPTA
AVSRRDPATPAEPVVRLDALATRLPDLEQRLEETCTDAFLVLRGSEVLAEYYRAGFAPDDRHLLMSVSKS
LCGTVVGALIDEGRIDPAQPVTEYVPELAGSVYDGPSVLQVLDMQISIDYNEDYVDPASEVQTHDRSAGW
RTRRDGDPADTYEFLTTLRGDGGTGEFQYCSANTDVLAWIVERVTGLRYVEALSTYLWAKLDADRDATI
TVDQTGFGFANGGVSCTARDLARVGRMMLDGGVAPGGRVVSQGWVESVLAGGSREAMTDEGFTSAF
PEGSYTRQWWCTGNERGNVSGIGIHGQNLWLDPRTDSVIVKLSSWPDPDTRHWHGLQSGILLDVSRAL
DAVLEHHHHHH

**Figure S7.** NylB DNA and Protein Sequences

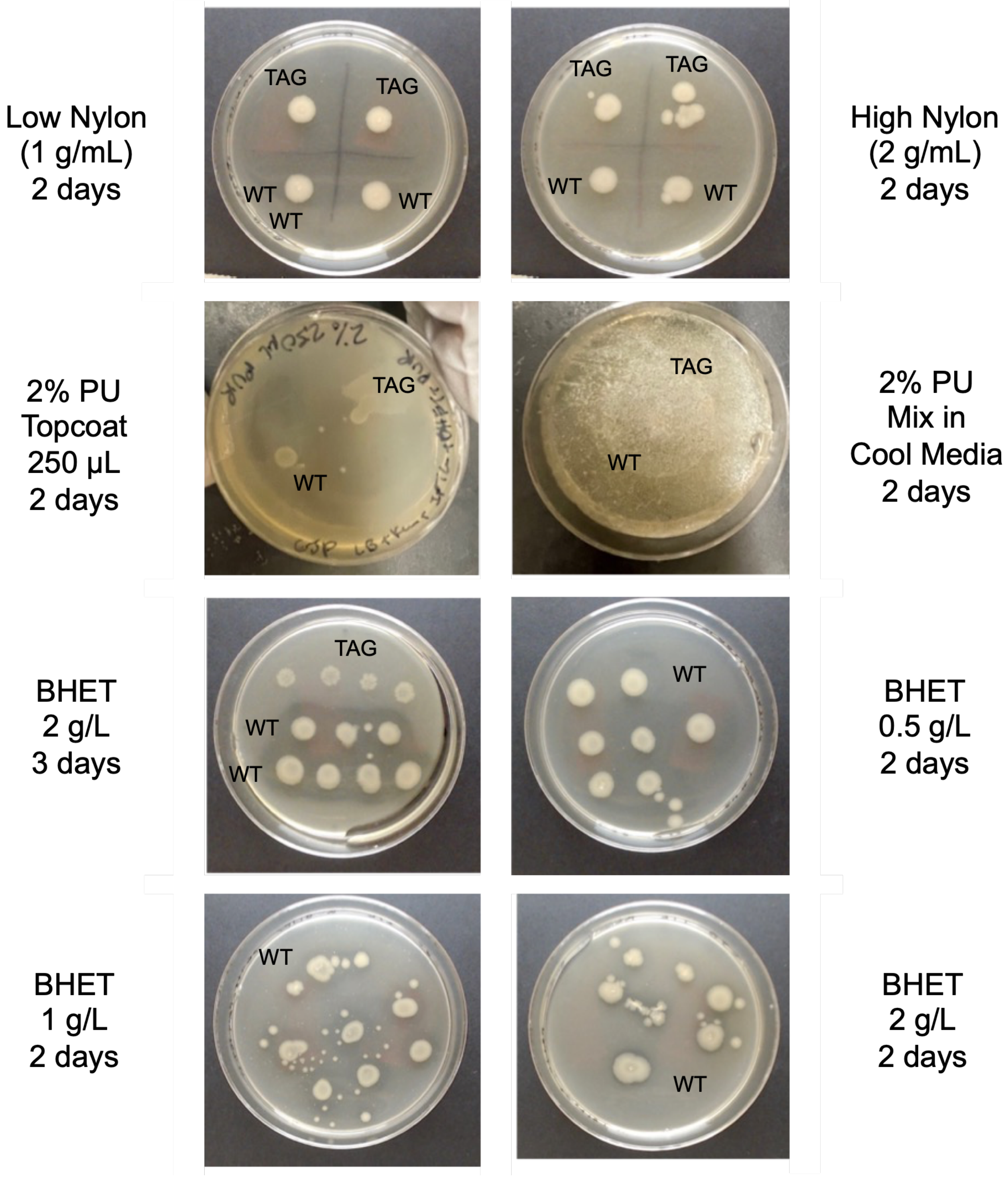


**Figure S8.** Representative Images of Polymer Plate Tests. Nylon, PU, and BHET preparations spotted with WT FAST PETase or TAG82 control plasmids and incubated for 2 or 3 days at 37 °C.

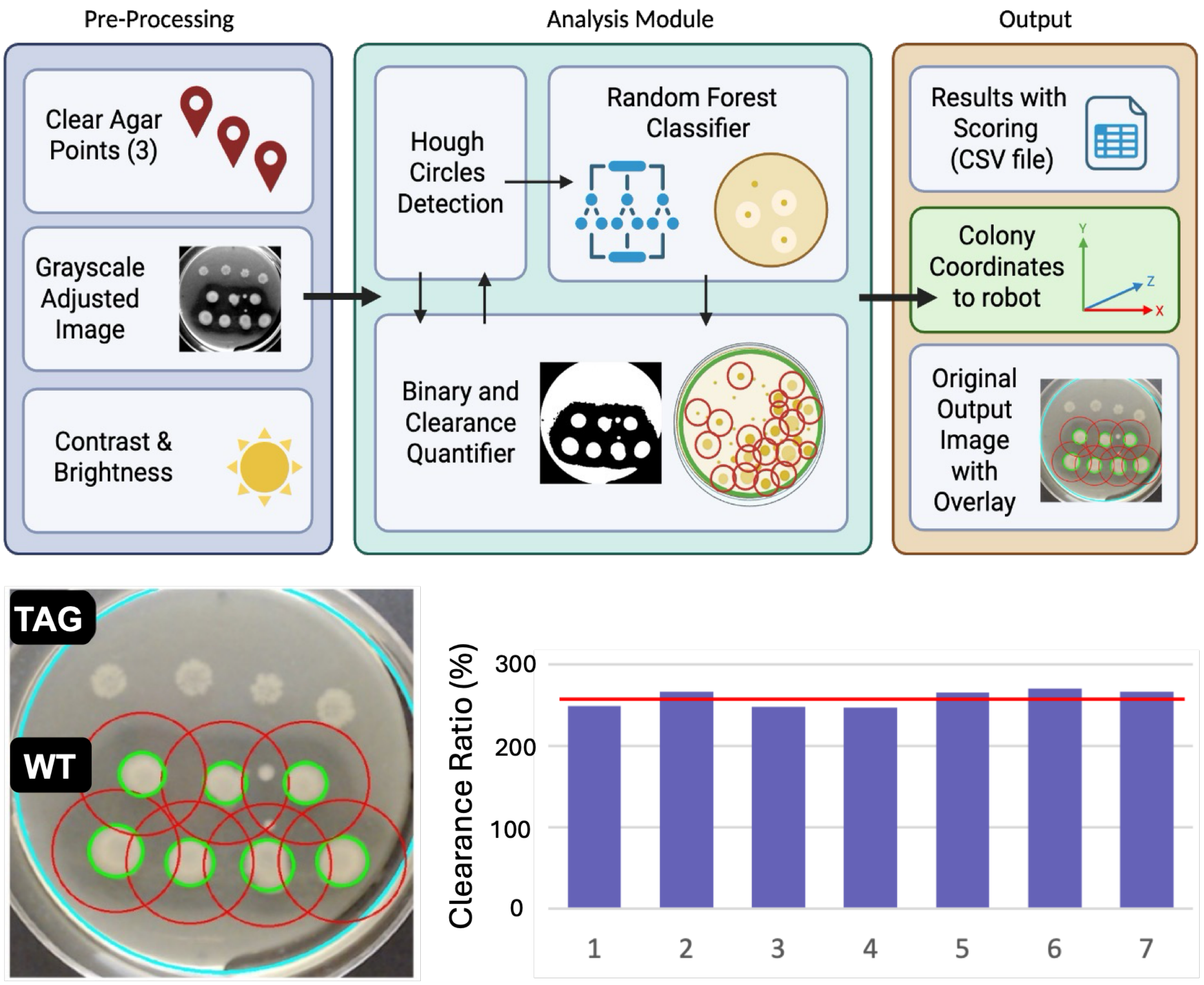


**Figure S9.** Plate Clearance Analysis. Top: Image analysis workflow with potential connection to colony picking robot. Bottom: Example plate analysis and quantification showing reproducibility.

Library 1 – BHET (left), PU (right)

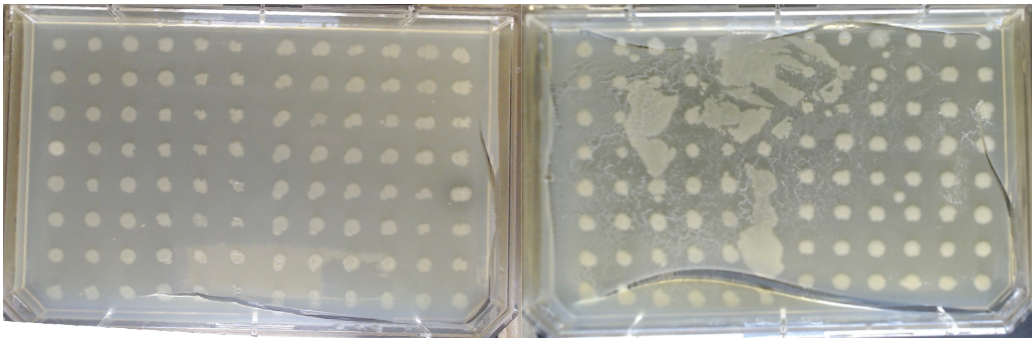

Library 2 – BHET (left), PU (right)

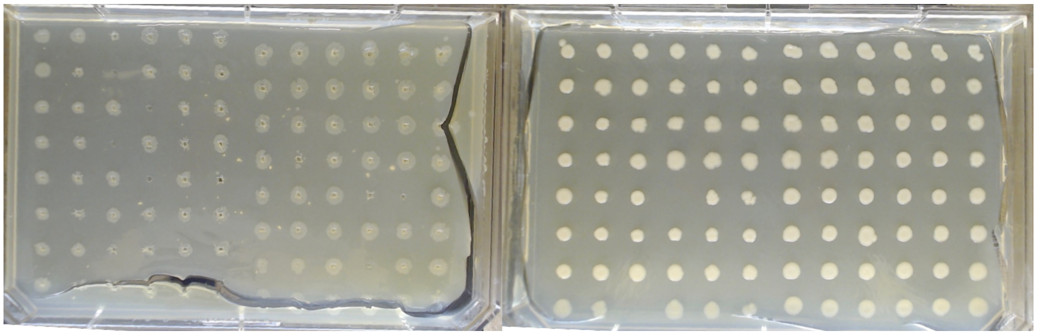

Library 3 – BHET (left), PU (right)

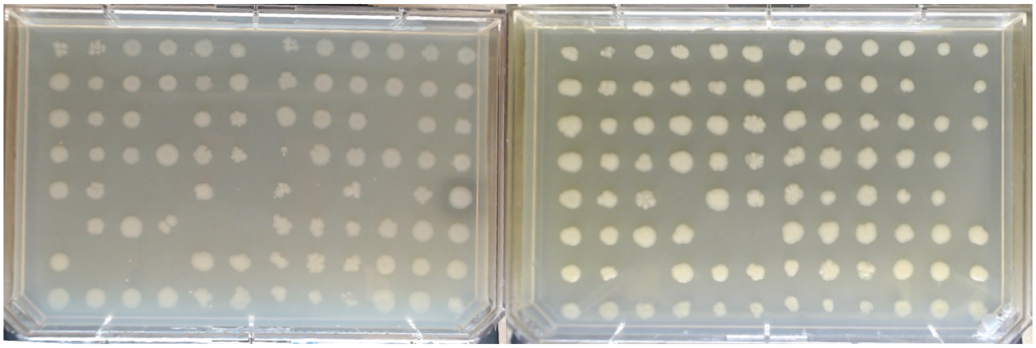

Library 4 – BHET (left), PU (right)

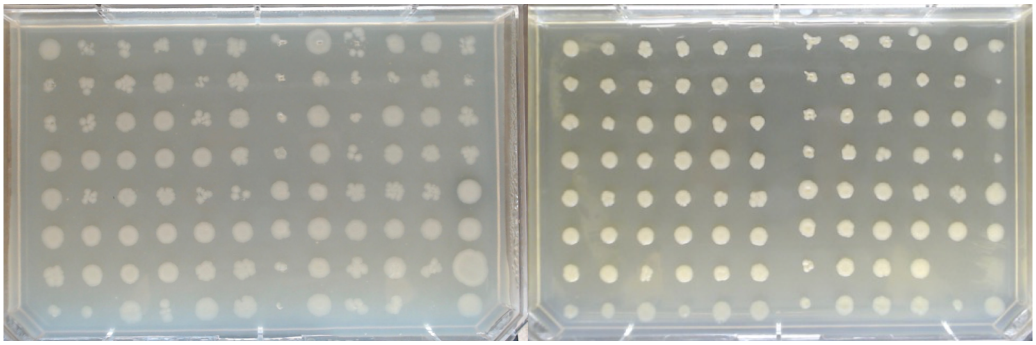

**Figure S10a.** Mutant Library Plate Images. Plates imaged after 2 days of growth at 37 °C.

Library 5 – BHET (left), PU (right)

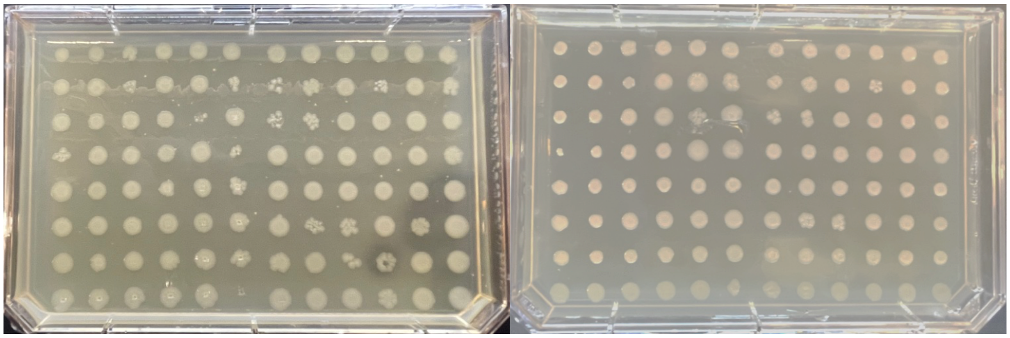

Library 6 – BHET (left), PU (right)

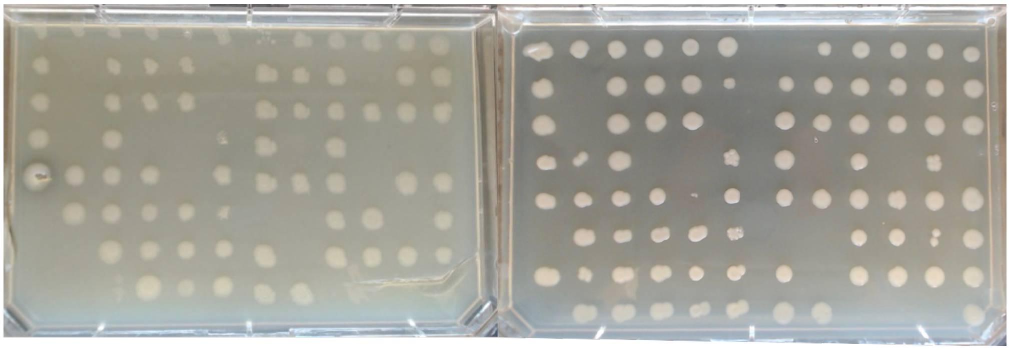

Library 7 – BHET (left), PU (right)

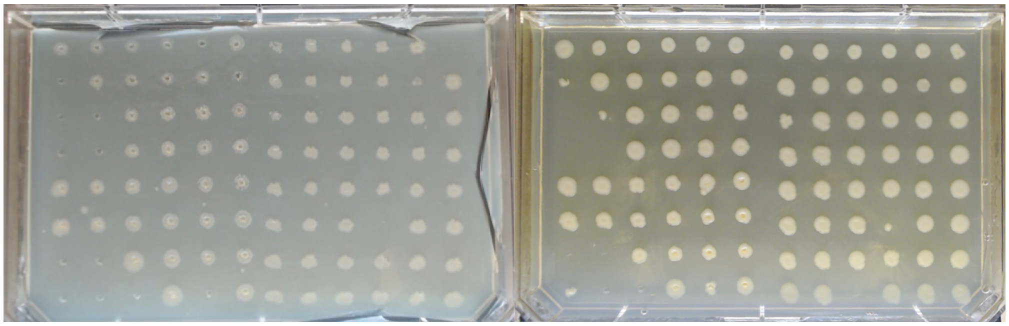

Library 8 – BHET (left), PU (right)

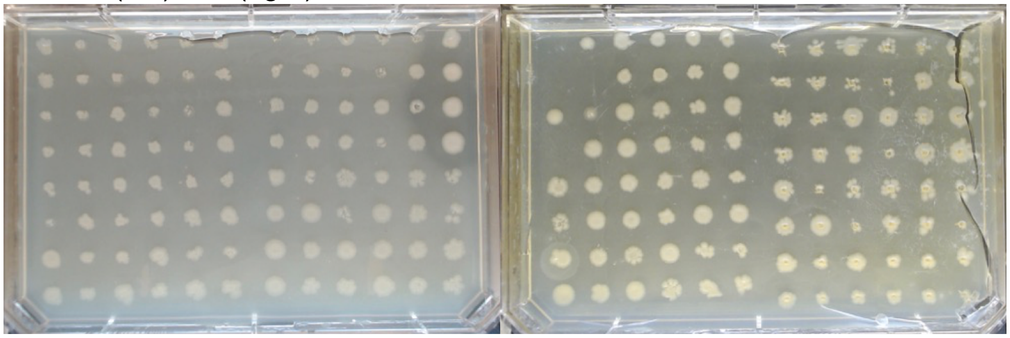

**Figure S10b.** Mutant Library Plate Images. Plates imaged after 2 days of growth at 37 °C.

Library 9 – BHET (left), PU (right)

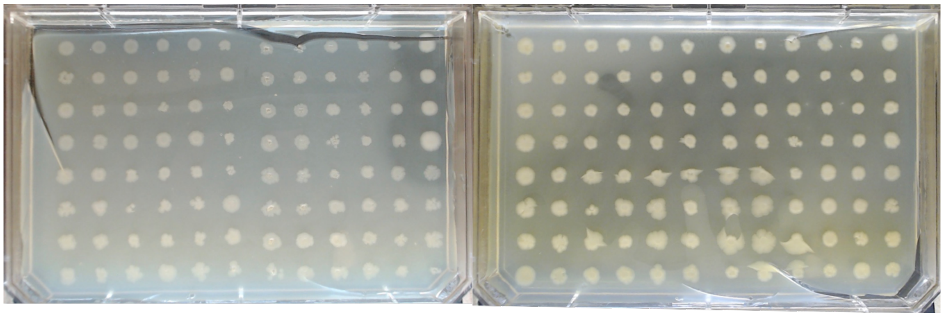

Library 10 – BHET (left), PU (right)

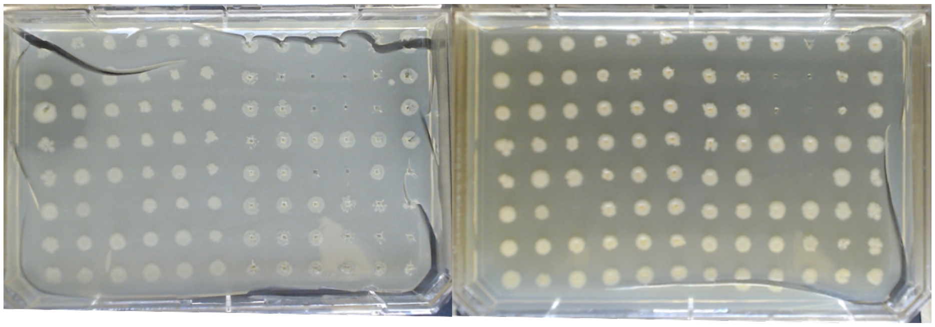

Library 11 – BHET (left), PU (right)

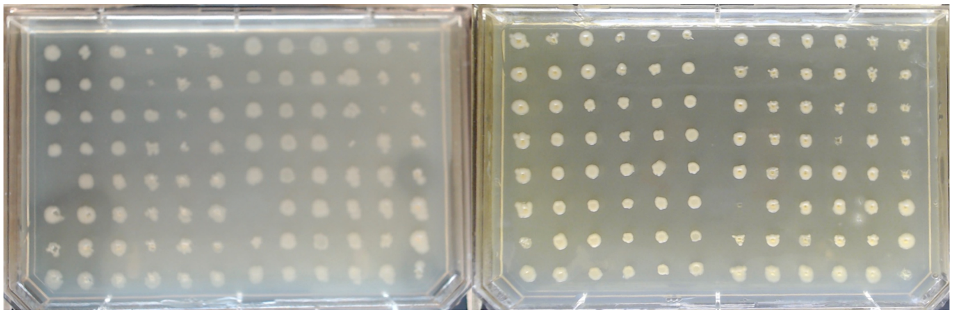

**Figure S10c.** Mutant Library Plate Images. Plates imaged after 2 days of growth at 37 °C.

**Sequences of Selected Mutants**

1. B1-D1 – W180G
2. B1-D11 – Y82V
3. B2-F7 – W154R, M156Q
4. B3-E6 – Y82N, W180G
5. U4-A7 – Y82I, W154A, M156K, W180V
6. U4-A8 – Y82R, M156V, W180Y
7. B4-A10 – Y82E, W154Z, M156D, W180G
8. B5-C5 – Y82W, W180V
9. B6-A5 – Y82Y, W180Y
10. U6-G7 – Y82W, W154R, M156A, W180G
11. U6-H7 – Y82L, W154V, M156V, W180G
12. U6-H8 – Y82E, W154Z, M156D, W180G

**Figure S11.** FAST PETase DNA and Protein Sequences

**P1-D1** –W180G

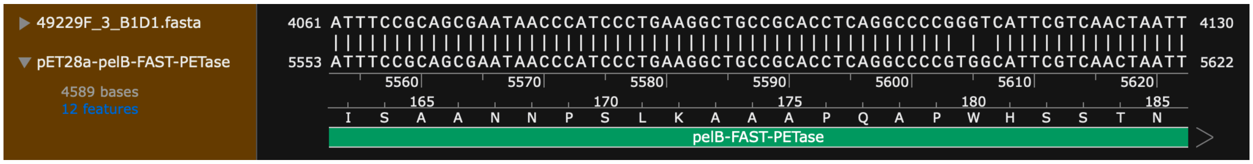


**P1-D11** –Y82V

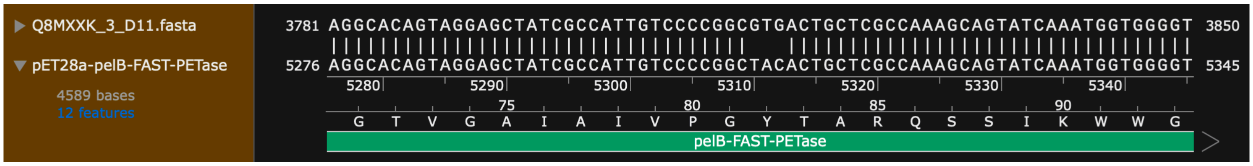


**P3-E6** – Y82N, W180G

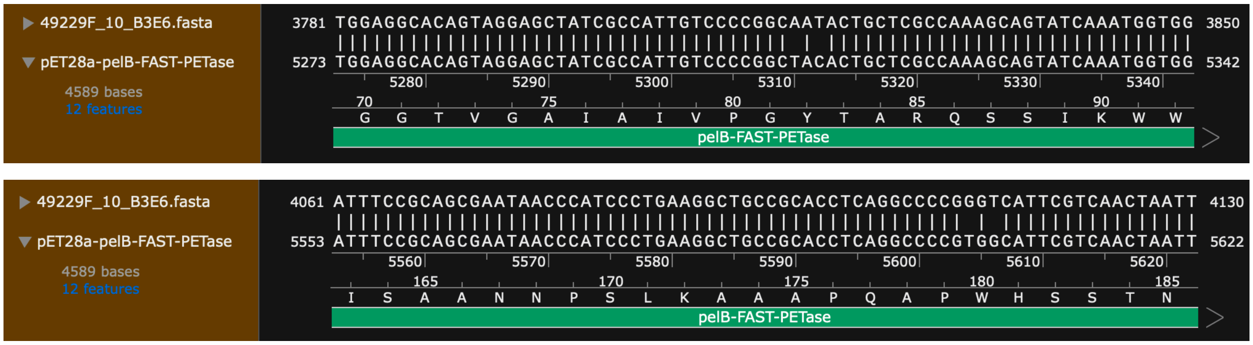


**P4-A7** – Y82I, W154A, M156K, W180V

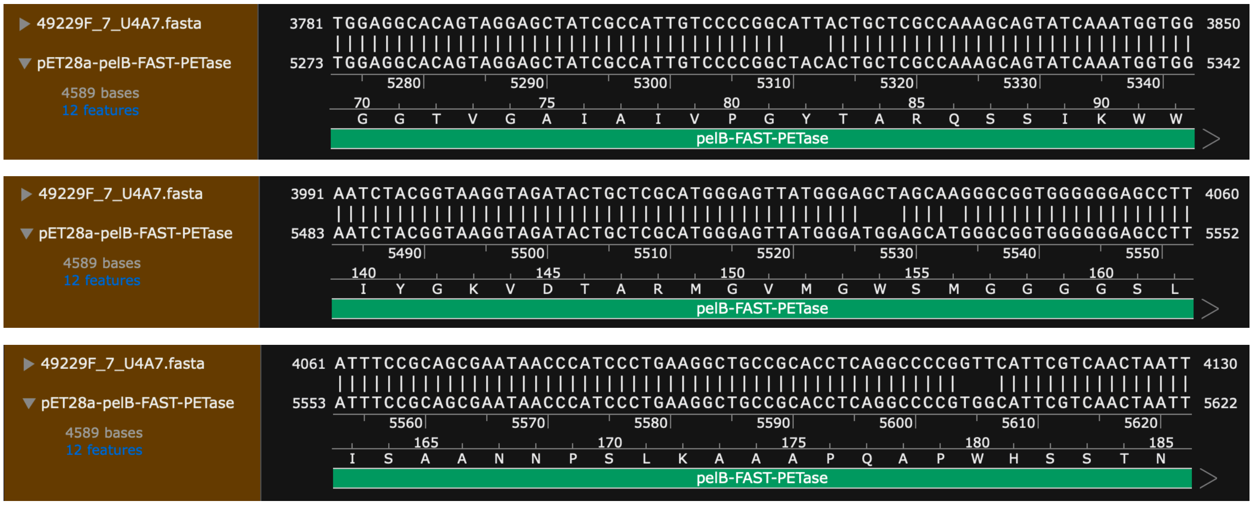


**Figure S12a.** Sequences of Selected FAST PETase Mutants.

**P4A10** – Y82E, W154Z, M156D, W180G

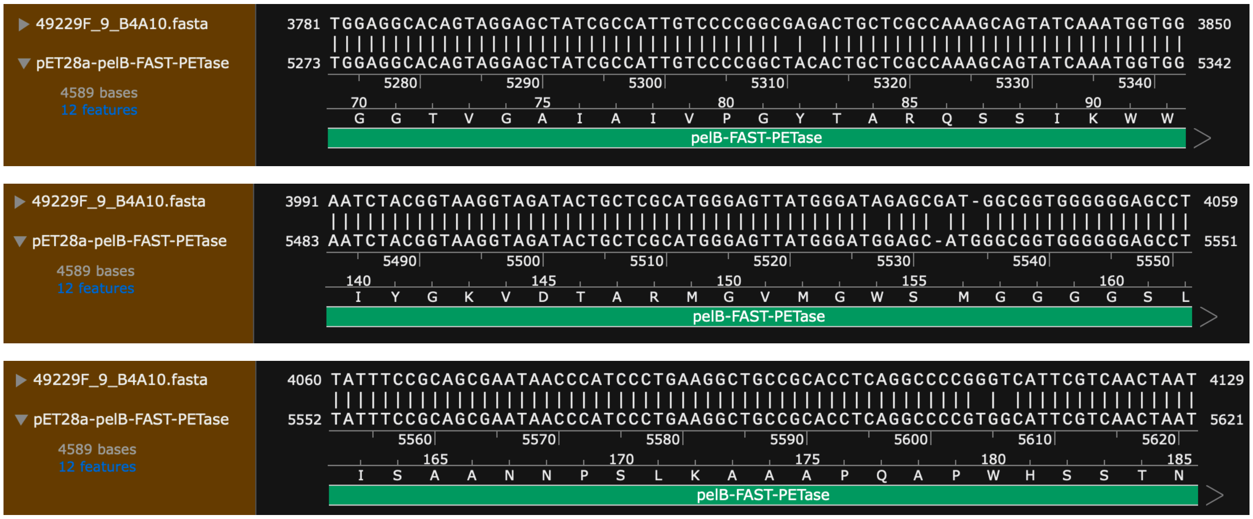


**P6-G7** – Y82W, W154R, M156A, W180G

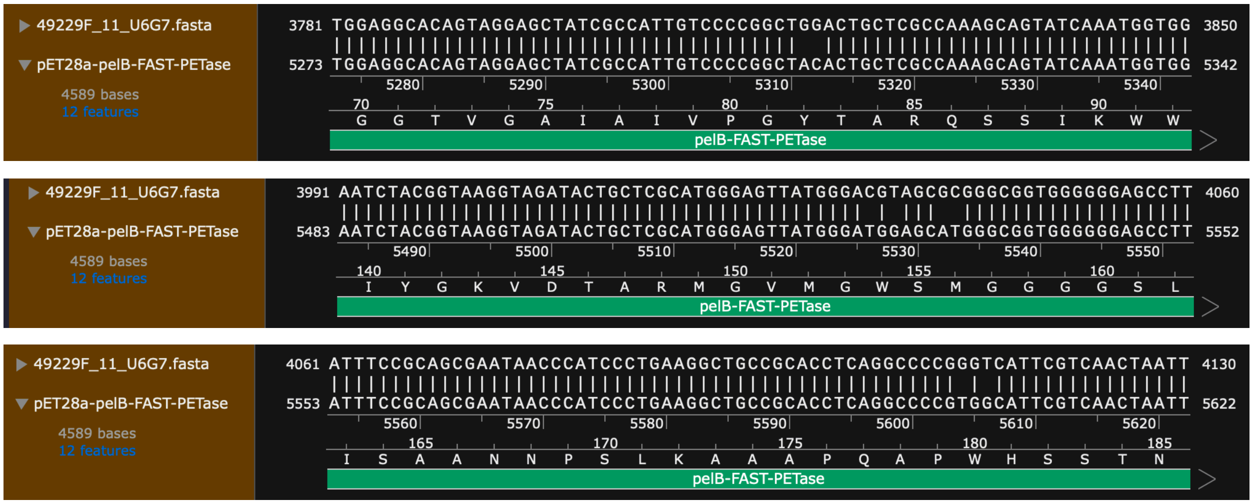


**Figure S12b.** Sequences of Selected FAST PETase Mutants.

**P6-H7** – Y82L, W154V, M156V, W180G

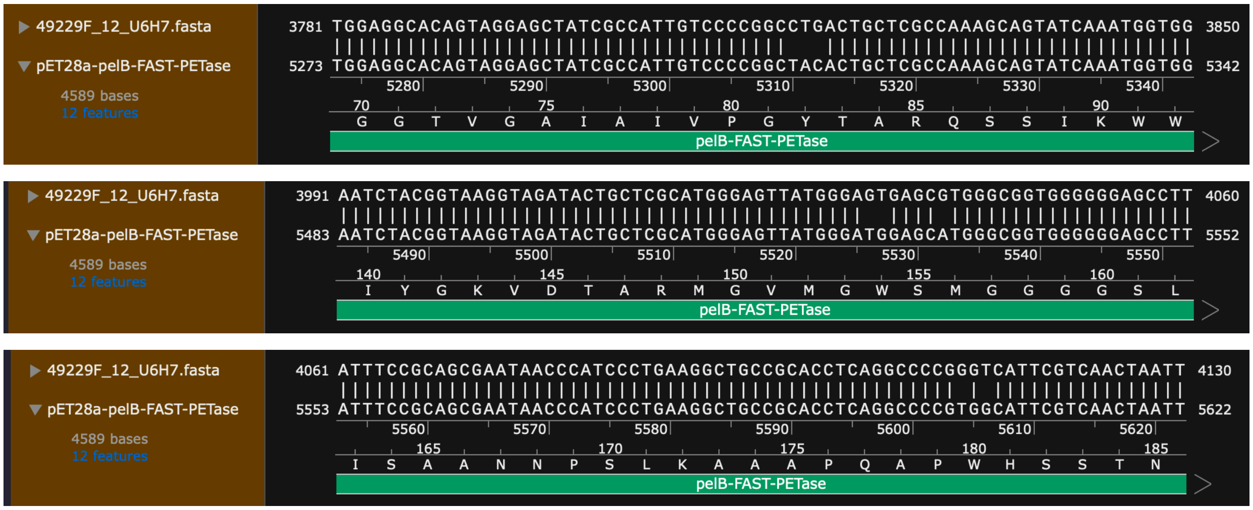


**P6-H8** –Y82E, W154Z, M156D, W180G

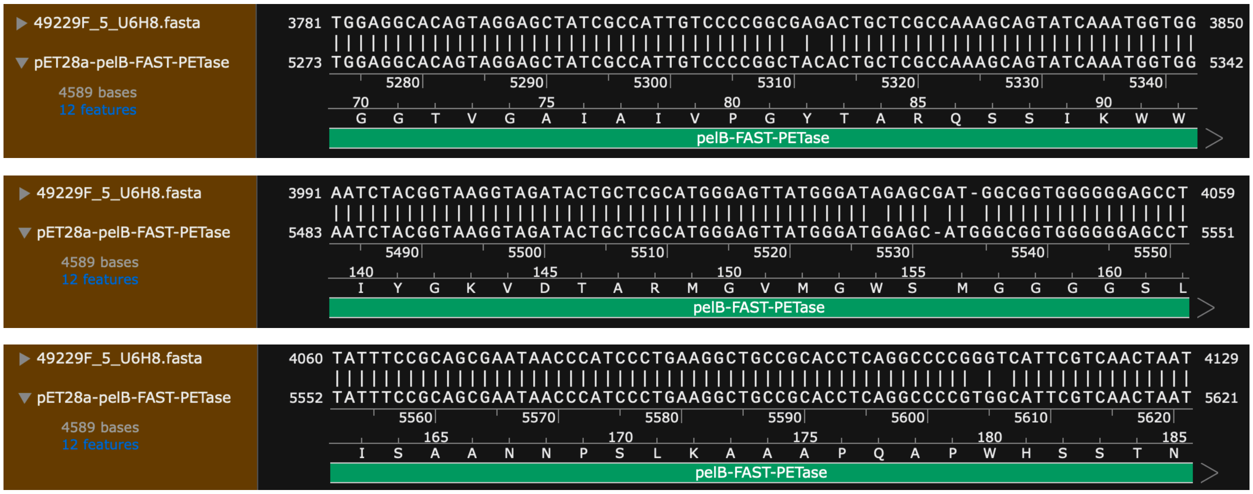


**Figure S12c.** Sequences of Selected FAST PETase Mutants.

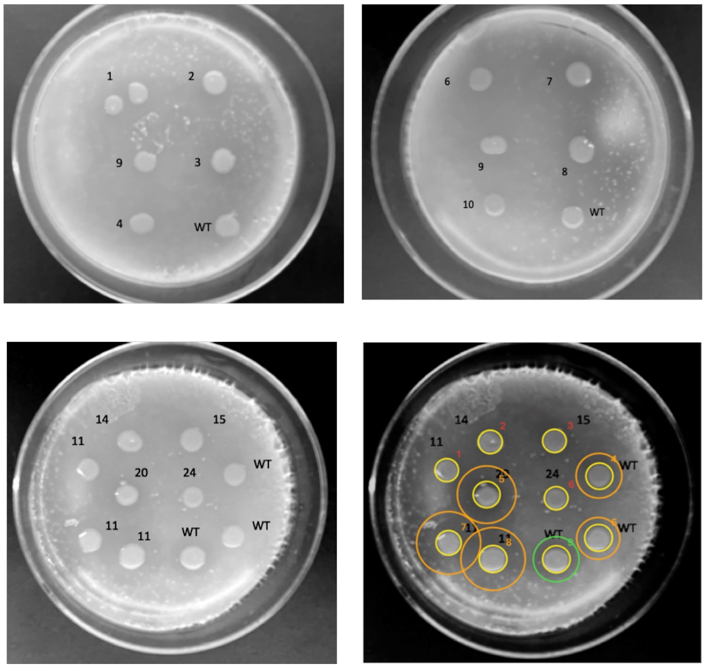

**Mutants**

1. U4A8 – Y82R, M156V, W180Y
2. U6H7 – Y82L, W154V, M156V, W180G
3. U4A7 – Y82I, W154A, M156K, W180V
4. B2F7 – W154R, M156Q
5. B5C5 – Y82W, W180V
6. B4A10 – Y82E, W154Z, M156D, W180G
7. U6G7 – Y82W, W154R, M156A, W180G
8. B3E6 – Y82N, W180G
9. B1D1 – W180G
10. U6H8 – Y82E, W154Z, M156D, W180G
11. B1D11 – Y82V
14. B10C1 – WT, possible mixture
15. B11F2 – Failed sequencing, possible mixture
20. B6A5 – W180Y
24. B5G10 – Y82L, W154H, W180T

**Figure S13.** PU Clearance Activity of Selected PETase Mutants. Images of colony growth and clearance on PU top-coated plate. **Inset:** PU clearance circle analysis reported in **Figure 7C**.

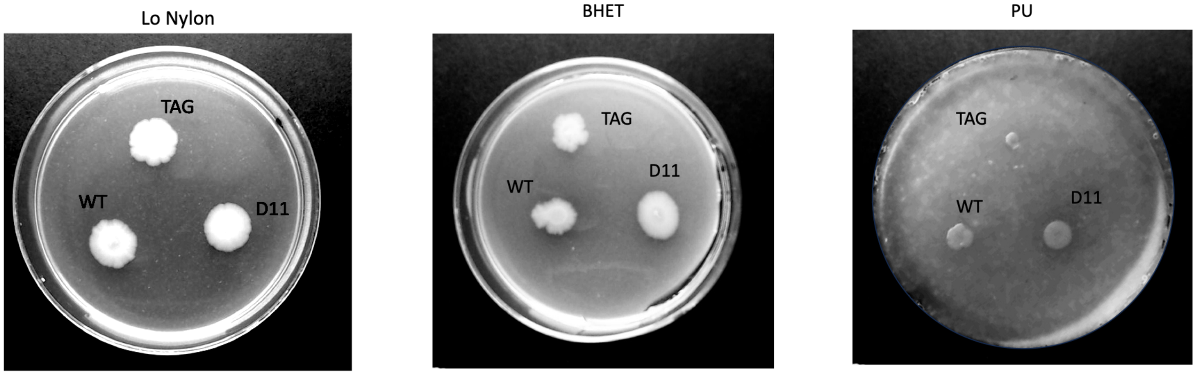


**Figure S14.** Y82V Mutant Does Not Clear Nylon. Images of colony growth and clearance on nylon- or BHET-embedded plates and PU top-coated plate. D11 mutant = Y82V.

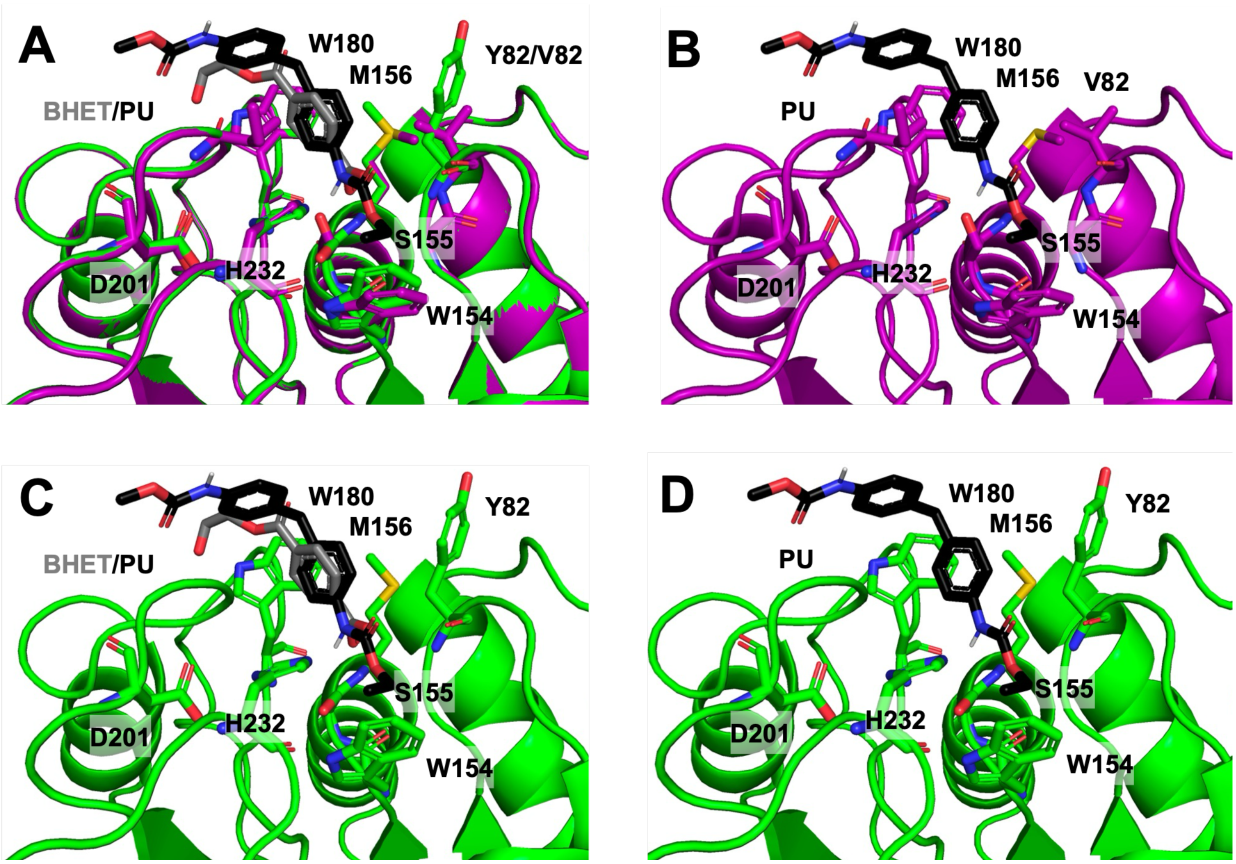


**Figure S15.** Sequences of Selected FAST PETase Mutants. **A**. Overlay of Y82V mutant after sculpting in PyMOL with PU substrate docked in the active site and FAST PETase structure 9LMV with BHET substrate from PDB ID 9LMV. **B.** Model of the Y82V with PU substrate docked in the active site. **C**. FAST PETase structure 9LMV with BHET substrate and PU substrate docked in the active site shown to highlight relative substrate positioning. **D**. FAST PETase structure 9LMV with PU substrate docked in the active site to show clash with M156.

**Table S1.** PETase Sequences from Wu et al. 2025

| | Residues | | | | | | | | |
|---|---|---|---|---|---|---|---|---|---|
| **PETase** | **81** | **82** | **154** | **155** | **156** | **180** | **201** | **203** | **232** |
| **APET-23** | G | A | H | S | F | I | D | G | H |
| **APET-24** | G | T | H | S | Q | G | D | V | H |
| **APET-25** | G | T | H | S | Q | G | D | I | H |
| **APET-26** | G | T | H | S | Q | G | D | I | H |
| **APET-27** | G | T | H | S | Q | Y | D | I | H |
| **APET-28** | G | T | H | S | Q | F | D | L | H |
| **LCC** | G | Y | H | S | M | W | D | V | H |
| **711** | G | Y | H | S | M | W | D | V | H |
| **607** | G | Y | H | S | M | W | D | V | H |
| **503** | G | Y | H | S | M | W | D | V | H |
| **606** | G | Y | H | S | M | W | D | V | H |
| **Thc_Cut1** | G | Y | H | S | M | W | D | I | H |
| **708** | G | Y | H | S | M | W | D | I | H |
| **717** | G | Y | H | S | M | W | D | I | H |
| **601** | G | F | H | S | M | W | D | I | H |
| **504** | G | F | H | S | M | W | D | I | H |
| **APET-15** | G | F | H | S | M | W | D | I | H |
| **APET-20** | G | F | H | S | M | W | D | I | H |
| **611** | G | F | H | S | M | W | D | V | H |
| **602** | G | F | H | S | M | W | D | V | H |
| **APET-19** | G | F | H | S | M | W | D | V | H |
| **405** | G | F | H | S | M | W | D | V | H |
| **APET-5** | G | F | H | S | M | W | D | I | H |
| **APET-29** | G | Y | H | S | L | Y | D | I | H |
| **407** | G | Y | H | S | M | F | D | T | H |
| **APET-32** | G | F | F | S | M | W | D | L | H |
| **KbPETase** | G | F | W | S | M | W | D | V | H |
| **CaPETase** | G | F | W | S | M | W | D | V | H |
| **403** | G | Y | W | S | M | W | D | V | H |
| **IsPETase** | G | Y | W | S | M | W | D | I | H |
| **APET-17** | G | Y | W | S | M | W | D | I | H |
| **401** | G | Y | W | S | M | W | D | I | H |
| **406** | G | Y | W | S | Y | K | D | T | H |
| **409** | P | Y | W | S | M | H | D | T | H |
